# Evaluating institutional open access performance: Sensitivity analysis

**DOI:** 10.1101/2020.03.19.998542

**Authors:** Chun-Kai Huang, Cameron Neylon, Richard Hosking, Lucy Montgomery, Katie Wilson, Alkim Ozaygen, Chloe Brookes-Kenworthy

## Abstract

In the article “Evaluating institutional open access performance: Methodology, challenges and assessment” we develop the first comprehensive and reproducible workflow that integrates multiple bibliographic data sources for evaluating institutional open access (OA) performance. The major data sources include Web of Science, Scopus, Microsoft Academic, and Unpaywall. However, each of these databases continues to update, both actively and retrospectively. This implies the results produced by the proposed process are potentially sensitive to both the choice of data source and the versions of them used. In addition, there remain the issue relating to selection bias in sample size and margin of error. The current work shows that the levels of sensitivity relating to the above issues can be significant at the institutional level. Hence, the transparency and clear documentation of the choices made on data sources (and their versions) and cut-off boundaries are vital for reproducibility and verifiability.

## 1 Introduction

Policy and implementation planning requires reliable, robust and relevant evidence. Much of the policy development for research resource allocation and strategy, including that focussed on the shift towards open access, relies on specific datasets, with known issues. Despite this, relatively few of the sources of evidence for research performance and evaluation test their sensitivity to the choices of data source and processing.

The evaluation of open access performance offers a useful case study of these issues. Early efforts to provide evidence on the extent of open access involved intensive manual data collection and analysis processes (Matsubayashi et al., 2009; Björk et al., 2010; Laakso et al., 2011). These do not lend themselves to regular updates or to tracking the effects of interventions. Much of the need for this manual work was driven by the challenges of discovering, identifying, disambiguating and determining the open access status of individual research outputs.

In general terms, analysing some aspect of research outputs, requires two sources of information. Firstly, data that connects outputs to the unit of analysis (e.g., organisation, funder, discipline). In our case we are interested in universities as a unit of analysis. Information linking research organisations to their outputs has traditionally been provided by the two major proprietary data providers, Web of Science and Scopus. More recently new players such as Digital Science have entered this space alongside Google Scholar and Microsoft Academic that both provide free services. Some affiliation data for research outputs is also provided by publishers through DOI registration agencies (particularly Crossref) but this is too patchy to be useful at this point.

The second source of information required is data linking individual outputs to the measure of interest. In our case this is open access status. The lack of a large-scale and comprehensive data source on open access status was the main reason driving the labour intensive and manual processes underpinning previous work. Over the past few years a number of services have emerged, with Unpaywall from OurResearch providing the most commonly used data.

A further critical issue is the means by which a set of outputs is unambigously connected to the data about those outputs in the second data source. This requires either that there be a shared unique identifier or a disambiguation process. As Unpaywall focuses on information about objects identified by Crossref DOIs we use Crossref as a source for attributes, such as publication date, that we require across all the outputs we examine. In other work we have shown that there are significant differences in the publication date recorded across different bibliographic data sources (Huang et al., 2020a).

In Huang et al. (2020b) we propose and present the first comprehensive and reproducible workflow that integrates multiple data sources for evaluating university open access levels. However, more detailed study is required to understand the implications of the data workflow and how sensitive it is against various data sources and decisions made regarding the use of these data. This companion white paper aims to provide detailed analyses on how the results can be sensitive to the choice of data sources, and inclusion of universities due to level of confidence.

### 1.1 Fully specifying an analysis workflow

In Figure 1 we describe the generic workflow for analysis of research outputs described above and the specifics of how this applies to the analysis of open access performance by research organisations. To achieve full transparency the ideal would be a fully specified, and therefore reproducible, workflow with specific instances of input and output data identified. This is not straightforward.

**Figure 1:**
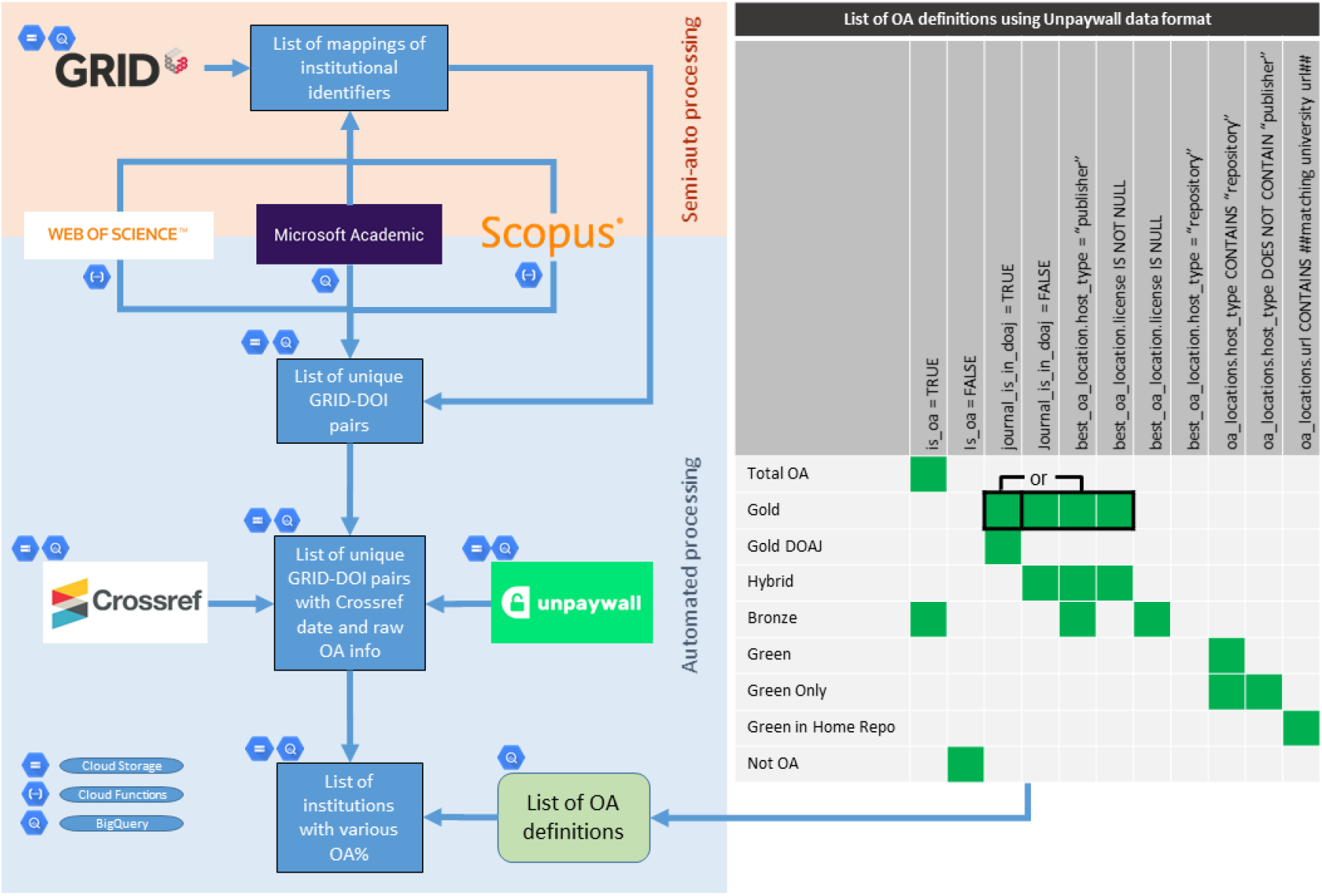
Workflow of data collection and mapping of open access definitions to Un-paywall metadata.

There are significant tensions between maintaining up-to-date data across multiple sources while simultaneously providing granular detail on the specific versions of those data used as inputs. Data collection is neither instantaneous nor synchronous. For example, for Microsoft Academic we can obtain timestamped data dumps which can be uniquely identified. However, for Web of Science and Scopus we use API calls which take time to complete. There are additional limitations due to licensing conditions on our ability to fully share the data obtained from some proprietary sources.

These conditions mean that it is not possible, either in theory or in practice, to provide a completely replicable workflow that would allow a third party to obtain exactly the same results. We can work towards a goal of fully describing as much of the workflow as possible, and provide usable snapshots of derived data products from as early in the pipeline as possible, but improving on this will be an ongoing process.

Note here that, in contrast to some of the analysis performed in the main article (Huang et al., 2020b), no filtering of the list of universities is applied in this article. However, a detailed analysis of the margins of error and sample size is given in Section 5.

### 1.2 The importance of sensitivity analysis

Because any such data analysis pipeline will involve making choices that cannot be made fully transparent, it is crucial that we provide an analysis of how large an effect those choices have on the results that we present. This includes the choices of data sources, issues of the timing of data collection, the statistical properties of output data and the likely robustness of specific metrics.

The remainder of this article is structured as follows. Section 2 compares the use of Web of Science, Scopus, Microsoft Academic and the full combined dataset. The comparisons are made at the country level, region level and over time. Section 3 examines the effects of using different Unpaywall versions. Lastly, Section 4 explores the relationship between sample size and margin of error related to the estimation of percentages of various open access categories.

## 2 Sensitivity analysis on the use Web of Science, Scopus and Microsoft Academic

In this section we explore in detail how the uses of the three different bibliographic indexing databases (in particular, Web of Science, Scopus and Microsoft Academic) affect the resulting open access scores at the institutional level. For details on the data collection process and our definitions of open access scores, please see the main article Huang et al., (2020b). The analyses are grouped into three subsections that examines differences at country level, region level and over time (per year), respectively.

### 2.1 Comparisons under groupings by country

Firstly, we reproduce the distributions of institutional OA scores (in terms of Total OA, Gold OA and Green OA) as in the main article (Huang et al., 2020b), grouped by country. This is labelled as Figure 2. Subsequently, we show the same display but restrict the underlying data to each of Microsoft Academic, Web of Science, and Scopus, respectively (Figures 3 to 5). In each of these figures, each dot represent the OA score of an university from the specific country, for 2017. The colour code for regions (as indicated in Figure 2) will be used throughout this article, where applicable.

**Figure 2:**
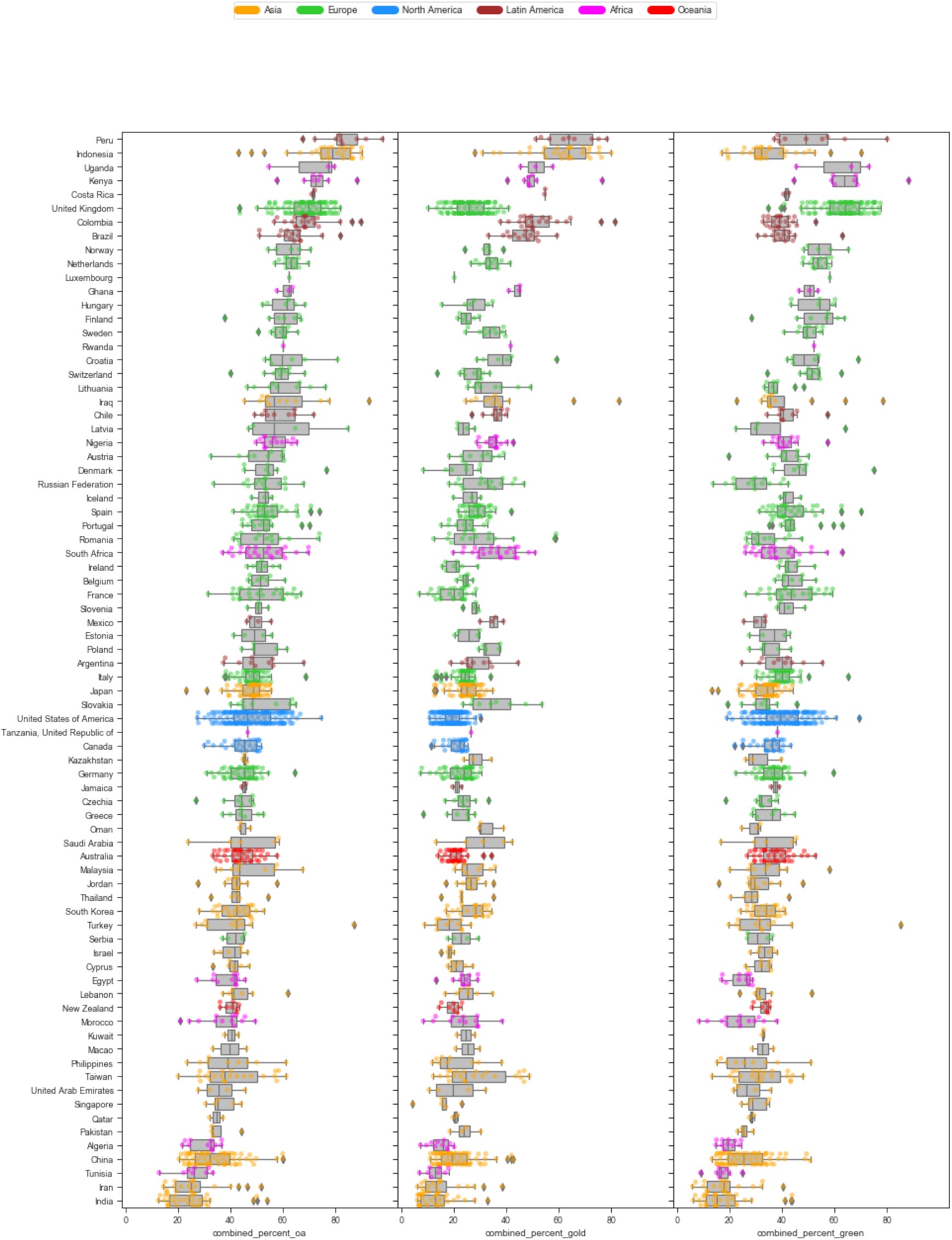
With the combined full dataset - 2017 percentages of Total OA, Gold OA and Green OA (left to right) grouped by countries.

**Figure 3:**
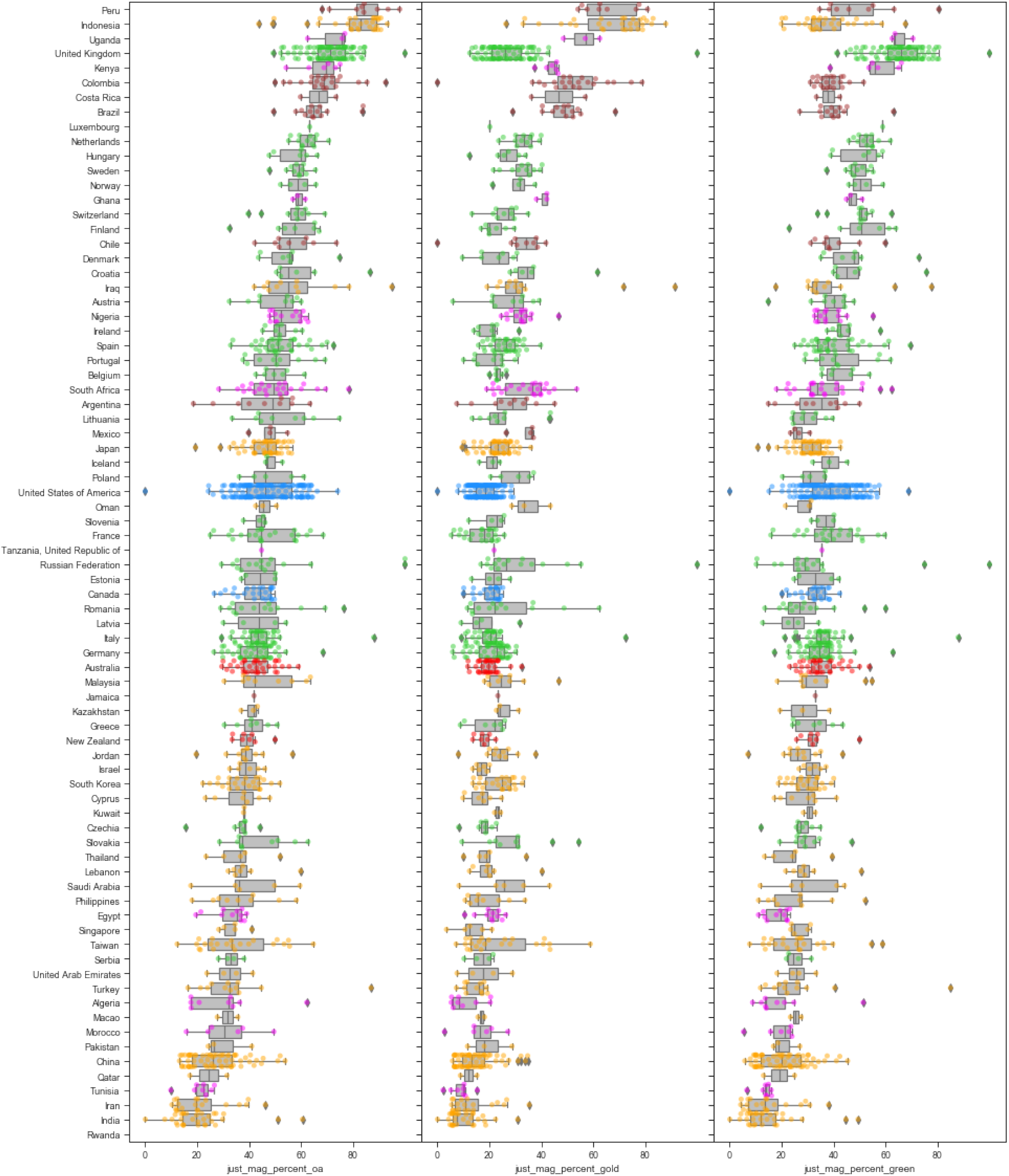
With only the Microsoft Academic dataset - 2017 percentages of Total OA, Gold OA and Green OA (left to right) grouped by countries.

**Figure 4:**
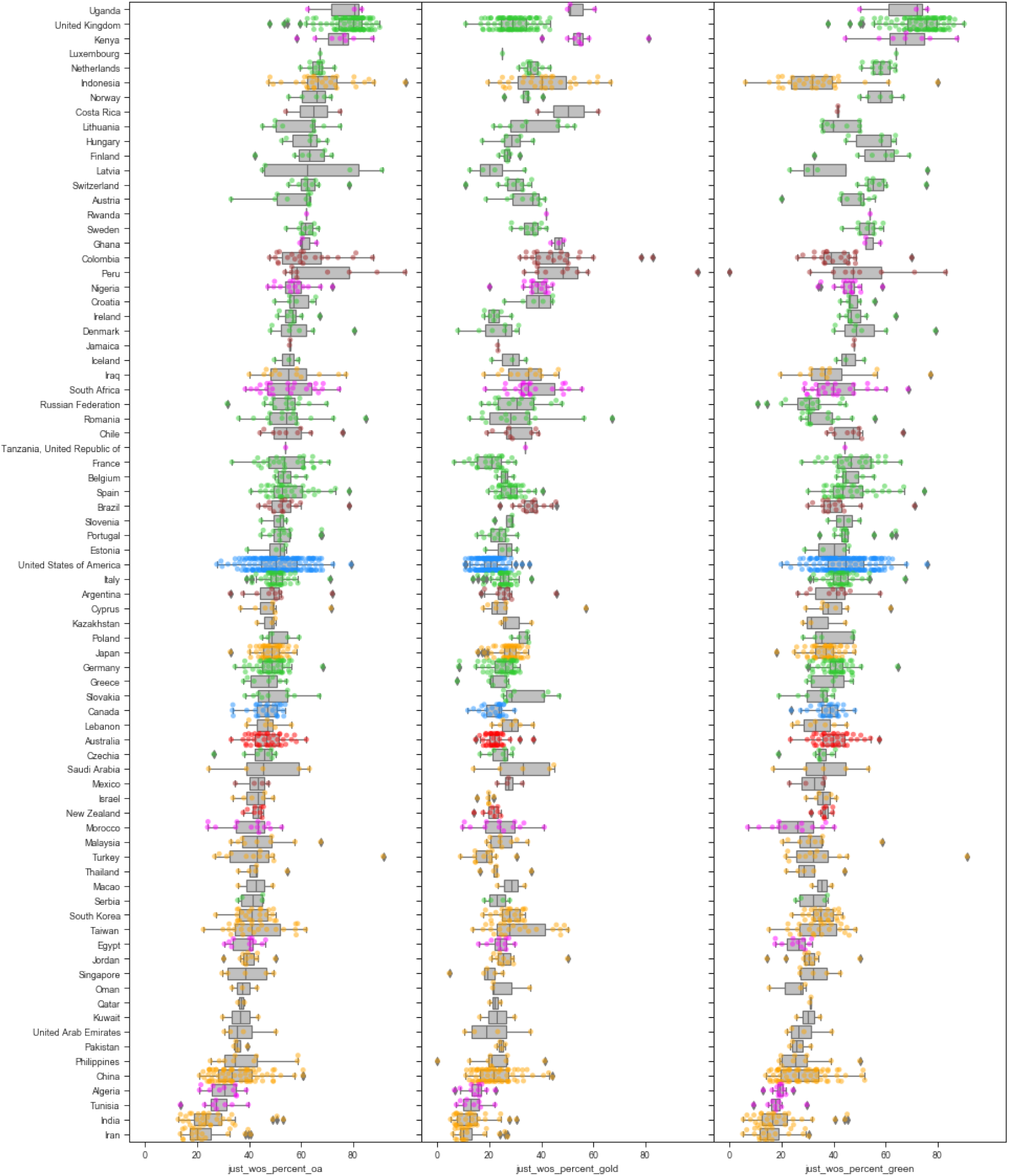
With only the Web of Science dataset - 2017 percentages of Total OA, Gold OA and Green OA (left to right) grouped by countries.

**Figure 5:**
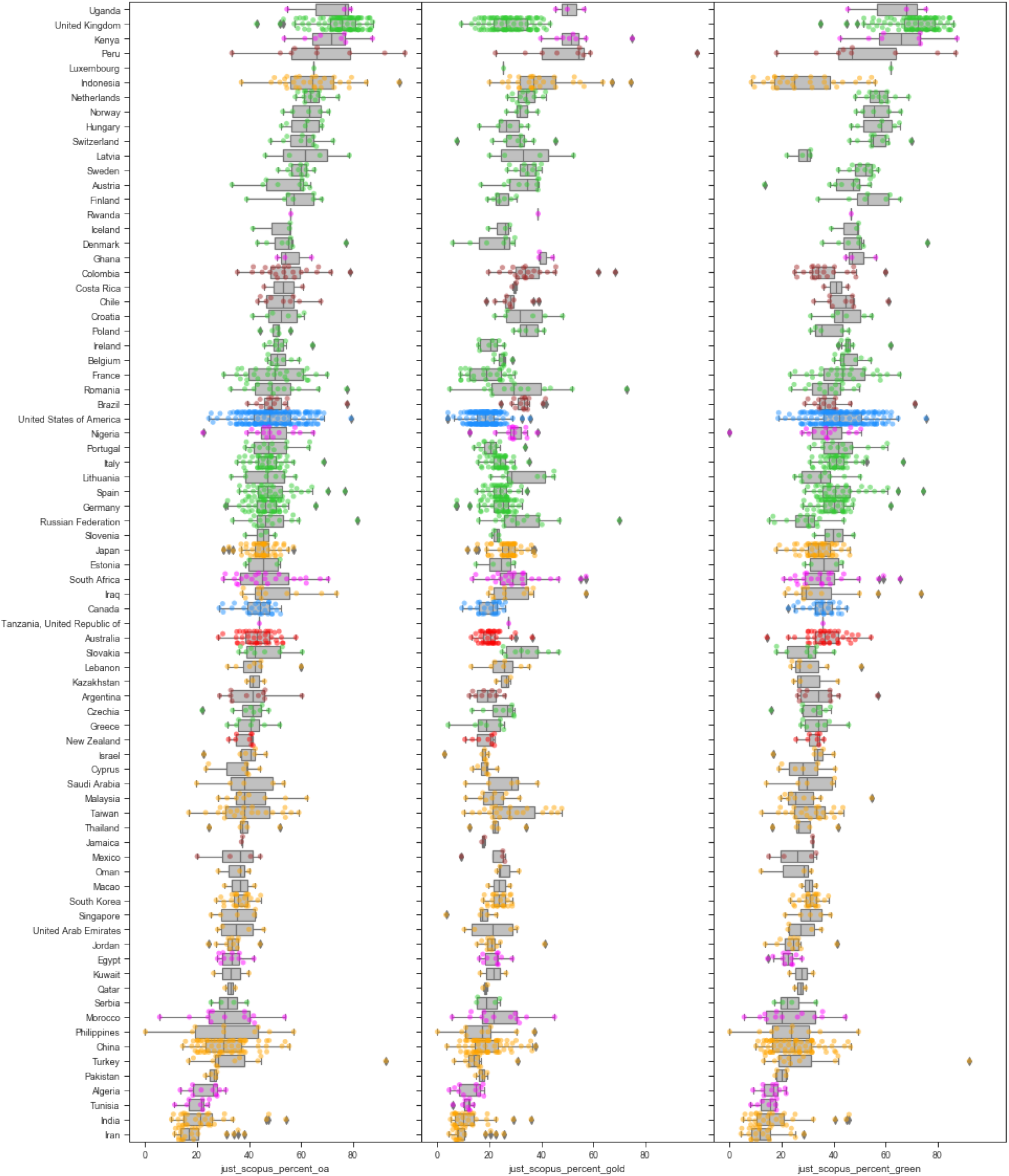
With only the Scopus dataset - 2017 percentages of Total OA, Gold OA and Green OA (left to right) grouped by countries.

In comparing Figures 2 to 5, we first note changes in the ordering of countries. In each figure, the order is determined by the median percentage of total open access (first column). These may be caused by the differences in coverage, as noted in Huang et al. (2020a). It is interesting to note the shifts of open access scores due to the choices of sources vary across countries. Not only do the median country open access scores shifts, the shapes of distributions of the various open access scores also change across datasets.

One of the most noticeable changes are that of the UK universities. The open access scores for UK universities remain mostly unchanged when the data is shifted from the full combined set to the one that only include Microsoft Academic records. However, the Green open access percentages seem to consistently shift upwards when using Web of Science or Scopus records only. This also results in higher total open access percentages for UK universities when only Web of Science or Scopus is used as the basis for DOI extraction.

To make the country level differences more clear, we now display the same graphs but for the open access percentage differences in Figures 6 to 11. Figures 6 to 8 shows the differences between the full dataset against each of Microsoft Academic, Web of Science, and Scopus. Figures 9 to 11 presents the differences amongst pairs of the three sources.

**Figure 6:**
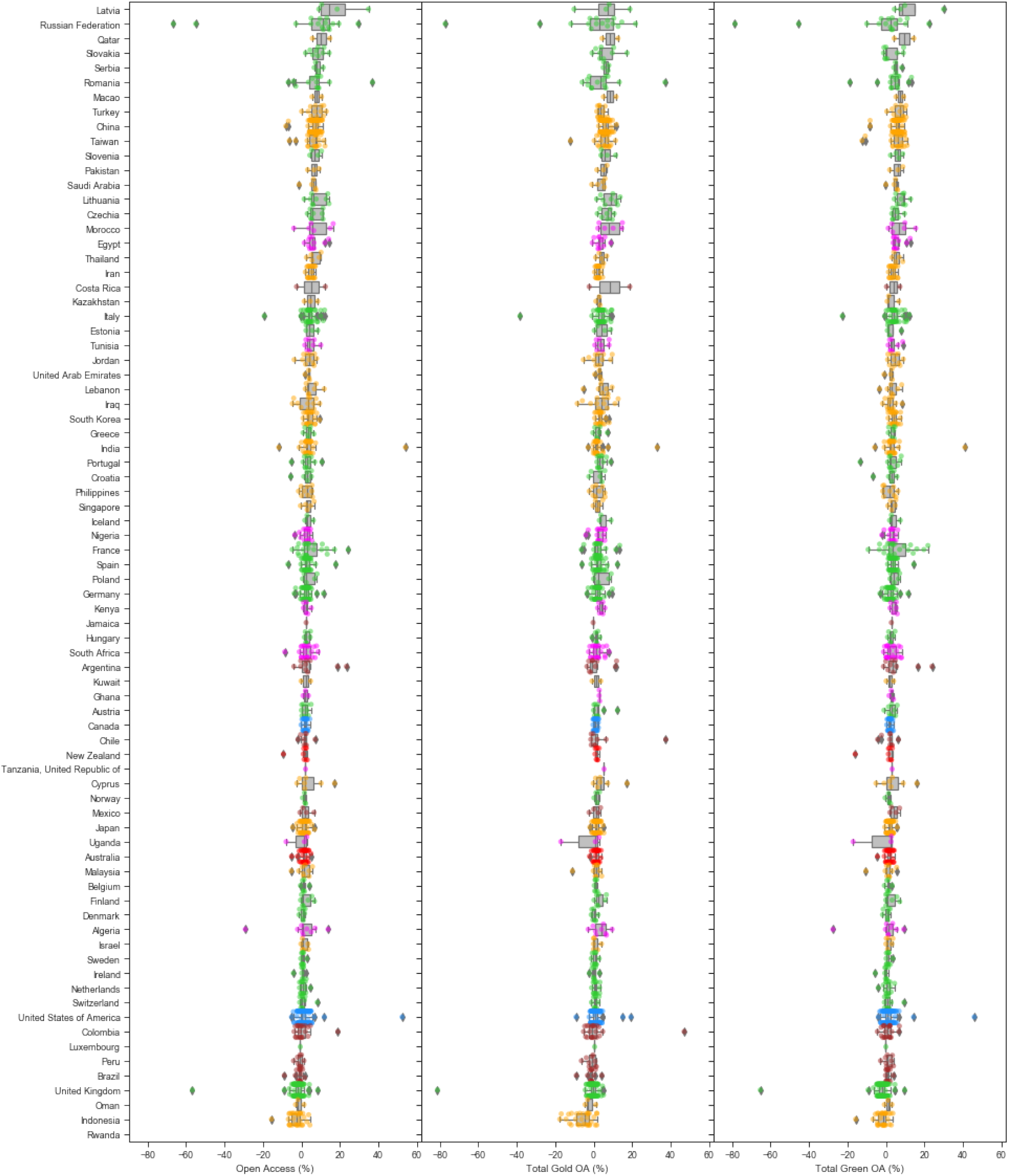
Differences in 2017 OA percentages for each institution between the full combined dataset and Microsoft Academic, grouped by country.

**Figure 7:**
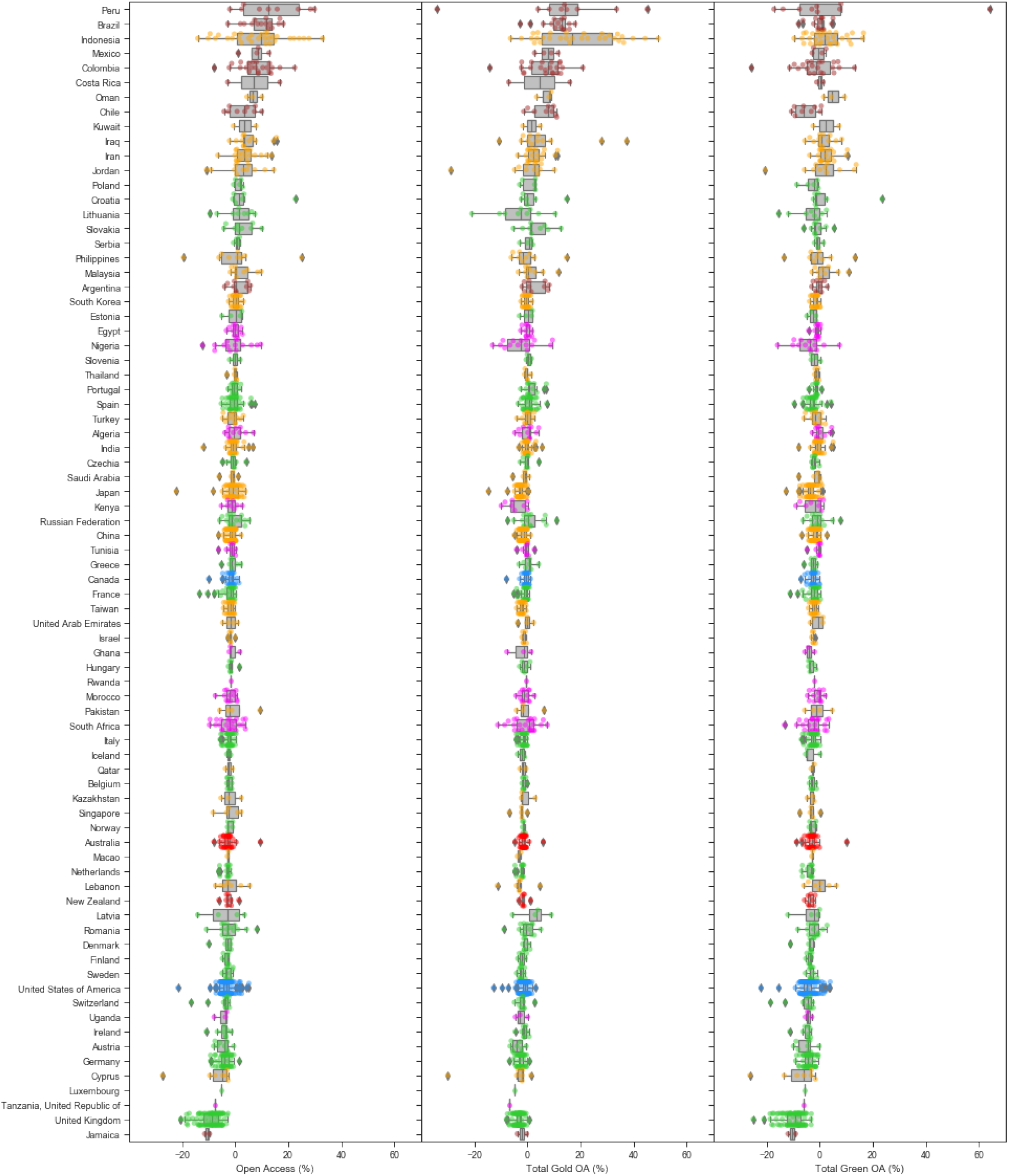
Differences in 2017 OA percentages for each institution between the full combined dataset and Web of Science, grouped by country.

**Figure 8:**
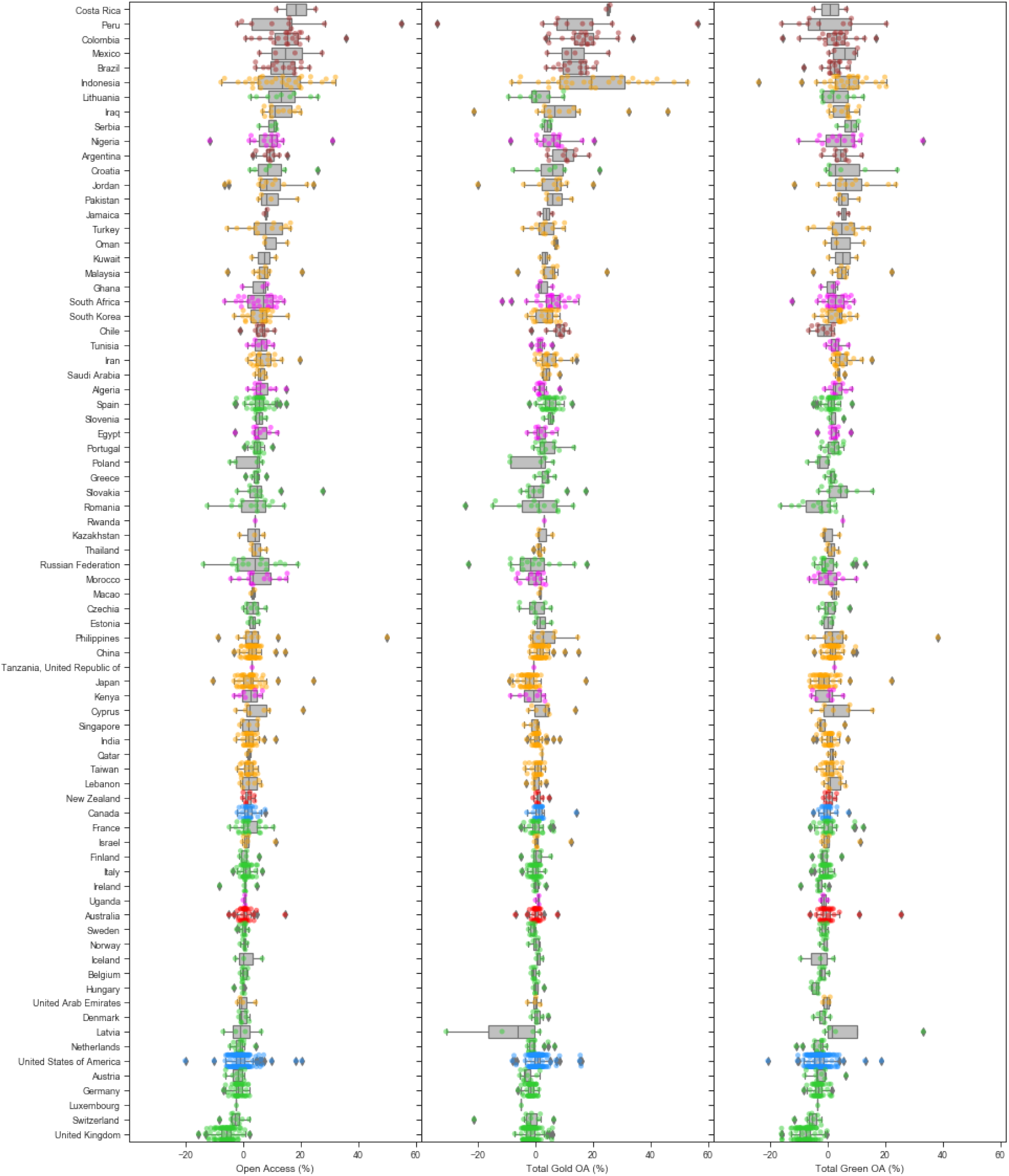
Differences in 2017 OA percentages for each institution between the full combined dataset and Scopus, grouped by country.

**Figure 9:**
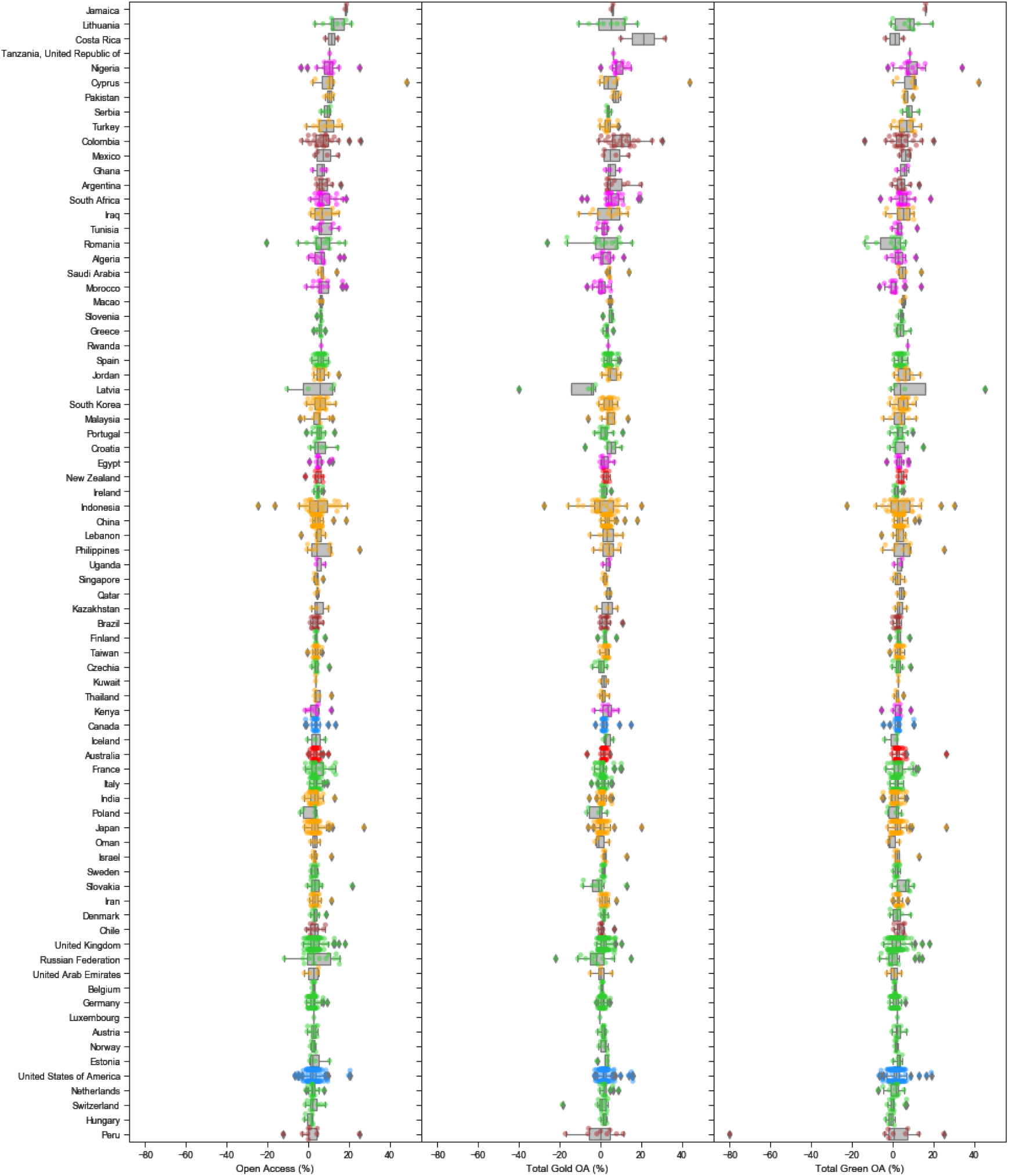
Differences in 2017 OA percentages for each institution between Web of Science and Scopus, grouped by country.

**Figure 10:**
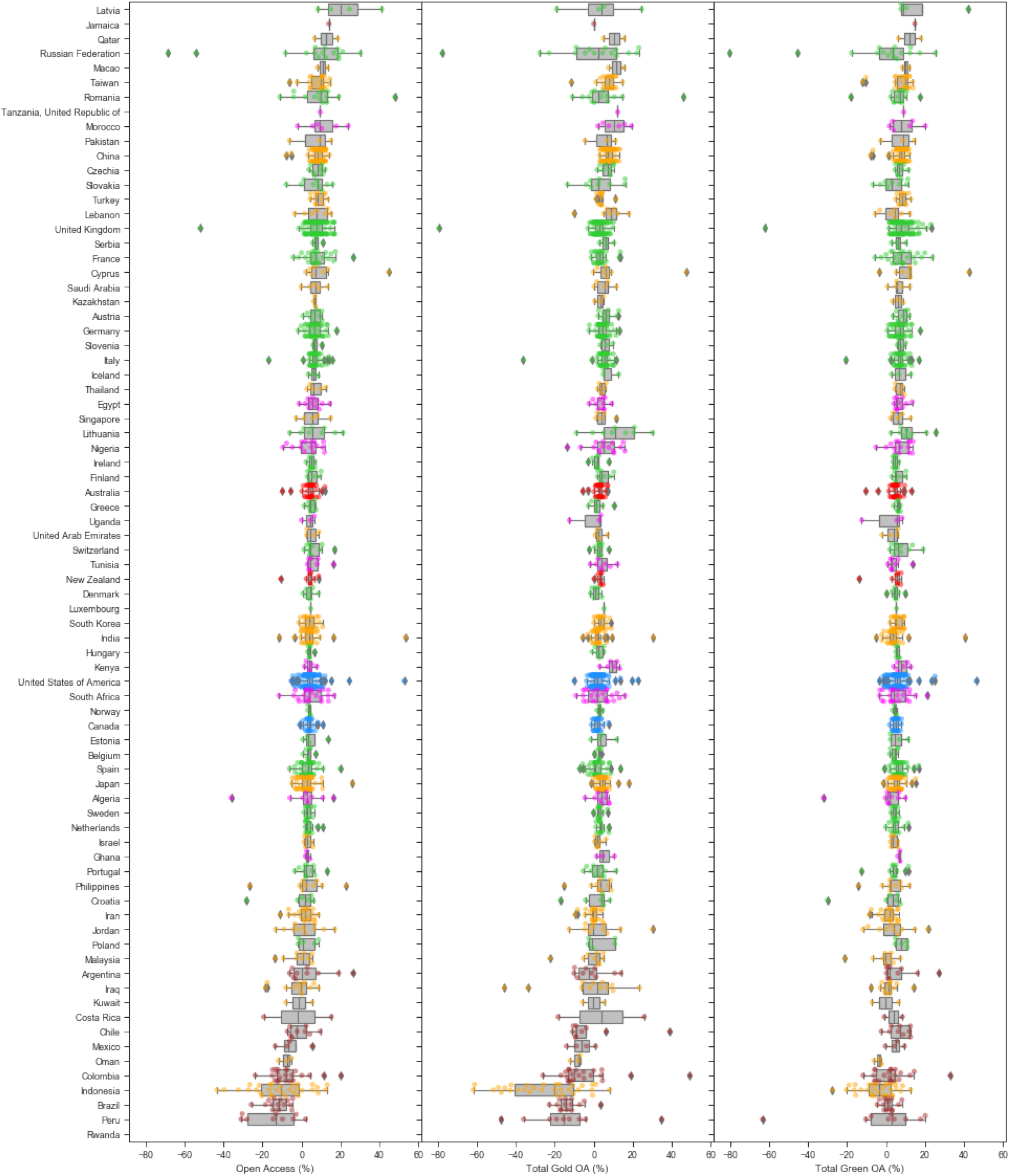
Differences in 2017 OA percentages for each institution between Web of Science and Microsoft Academic, grouped by country.

**Figure 11:**
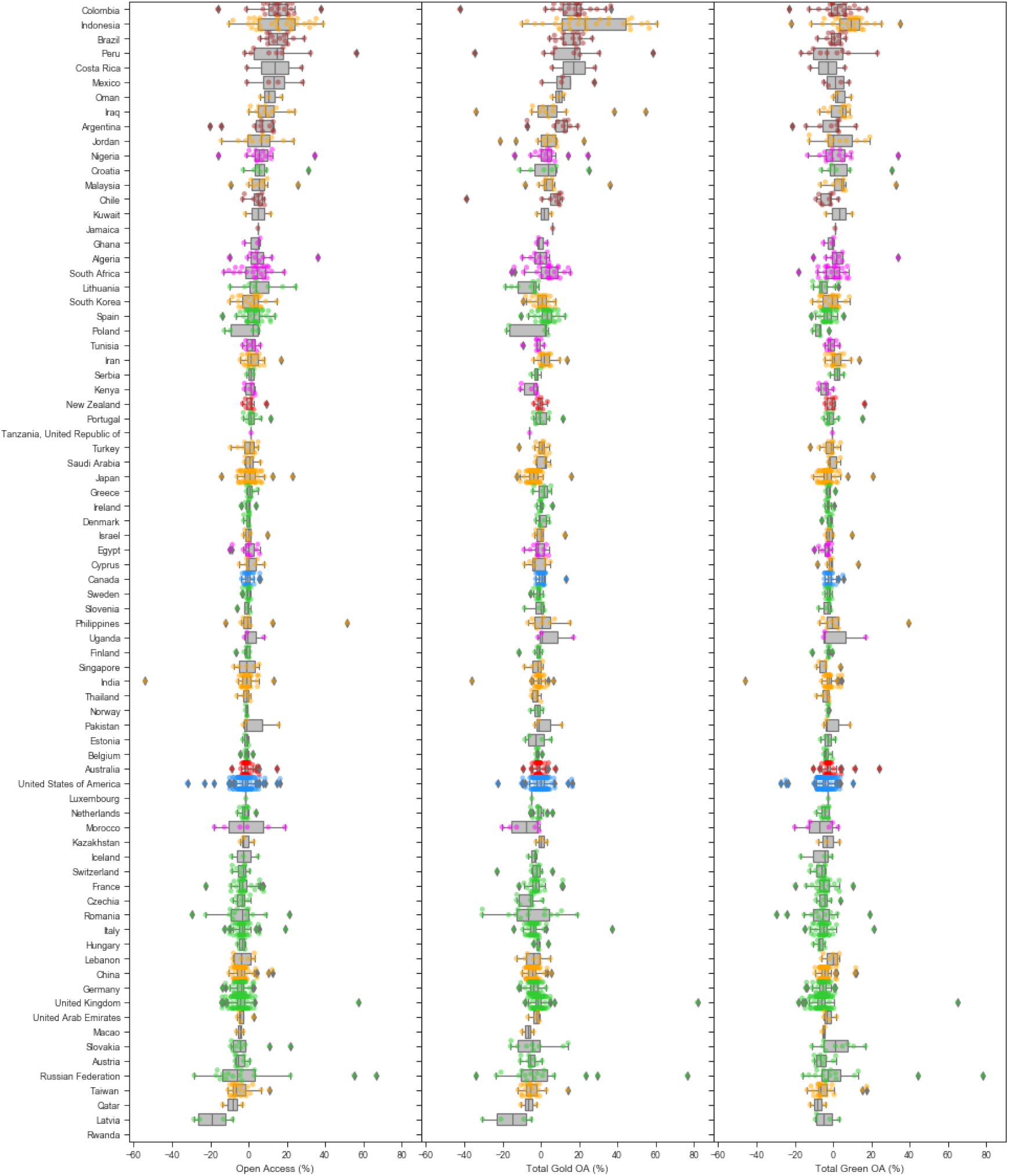
Differences in 2017 OA percentages for each institution between Microsoft Academic and Scopus, grouped by country.

In each of the Figures 6 to 11, the countries are ordered by the median differences in total open access percentages. For total open access, the shifts are mainly positive for universities in Brazil, Indonesia and Mexico, when the full combined dataset is used instead of Web of Science or Scopus only. The detailed effects on gold open access and green open access levels differ from that of total open acess. In particular, these universities’ gold open access percentages are higher (positive shifts), and green open access percentages lower (negative shifts), for the full combined dataset. In contrast, there are negative shifts for all three categories of open access relative to Web of Science and Scopus. The parallel comparison of the full combined dataset against Microsoft Academic shows substantially smaller differences, with most countries depicting a median difference around zero.

The comparisons between the three data sources show relatively less differences in results between Web of Science and Scopus, when contrasted against their differences in results against Microsoft Academic. These are indications that Microsoft Academic potentially have more comprehensive coverage of research outputs, especially for a select number of countries. This is consistent with what we have previously observed (Huang et al., 2020a).

### 2.2 Comparisons under groupings by region

As an alternative view of the above, we next group the universities into their respective regions and compares the resulting differences across these regions. Results are again for the year 2017. Figure 12 presents the various boxplots of the differences in total open access percentages (row 1), gold open access percentages (row 2) and green open access percentages (row 3) when contrasting the full combined dataset against each of Web of Science (column 1), Scopus (column 2) and Microsoft Academic (column 3). Figure 13 displays parallel visualisations for comparing open access percentage differences across any pairs of Web of Science, Scopus and Microsoft Academic.

**Figure 12:**
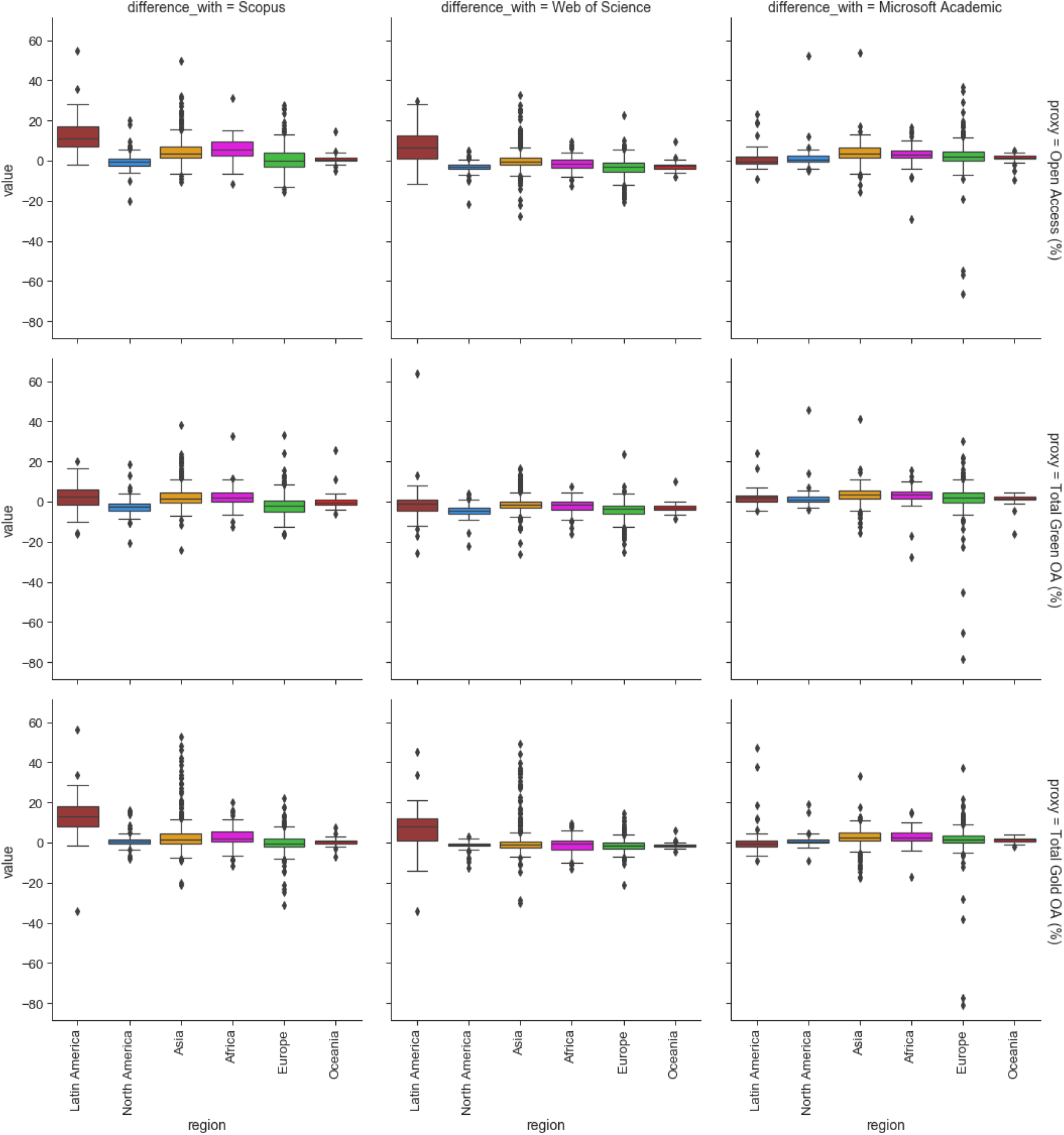
Differences in OA% when contrasting the full combined dataset again each of Web of Science, Scopus and Microsoft Academic.

**Figure 13:**
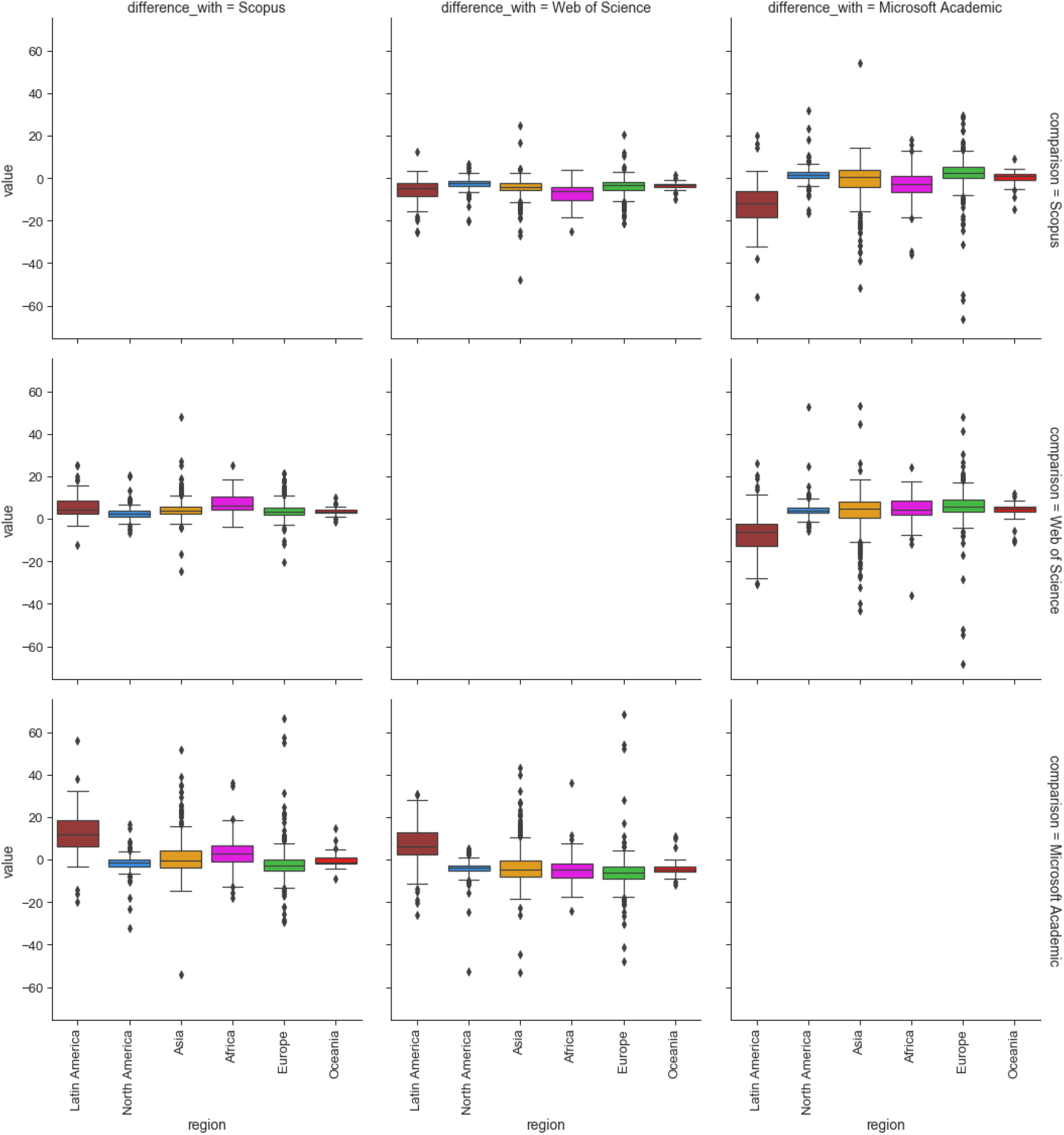
Differences in OA% when contrasting pairs of Web of Science, Scopus and Microsoft Academic.

The figures clearly depicts an advantage for Latin American universities when the full combined dataset, or Microsoft Academic, is used as opposed to using only Webs of Science or Scopus. The indication that Microsoft Academic produces a similar overall result to the full combined dataset is also re-emphasised in these graphs. However there remain large differences for specific universities. It is also worth noting the high number of outliers for Asian universities when contrasting the the combined set against Web of Science and Scopus, and similarly when comparing across pairs of the three data sources.

### 2.3 Comparisons of differences across years

This subsection explores how the differences in various OA% across different pairs of datasets vary across years. Readers should note that the abnormality observed for the year 2019 is mainly due to the time delay in data collection, particularly from Web of Science and Scopus. We leave 2019 in the visualisation to illustrate some of the limitations of our data pipeline. Figure 14 describes the differences in total open access percentages between the combined data set and each of the individual data sources. Corresponding figures for gold open access and green open access are presented in Figures 15 and 16.

**Figure 14:**
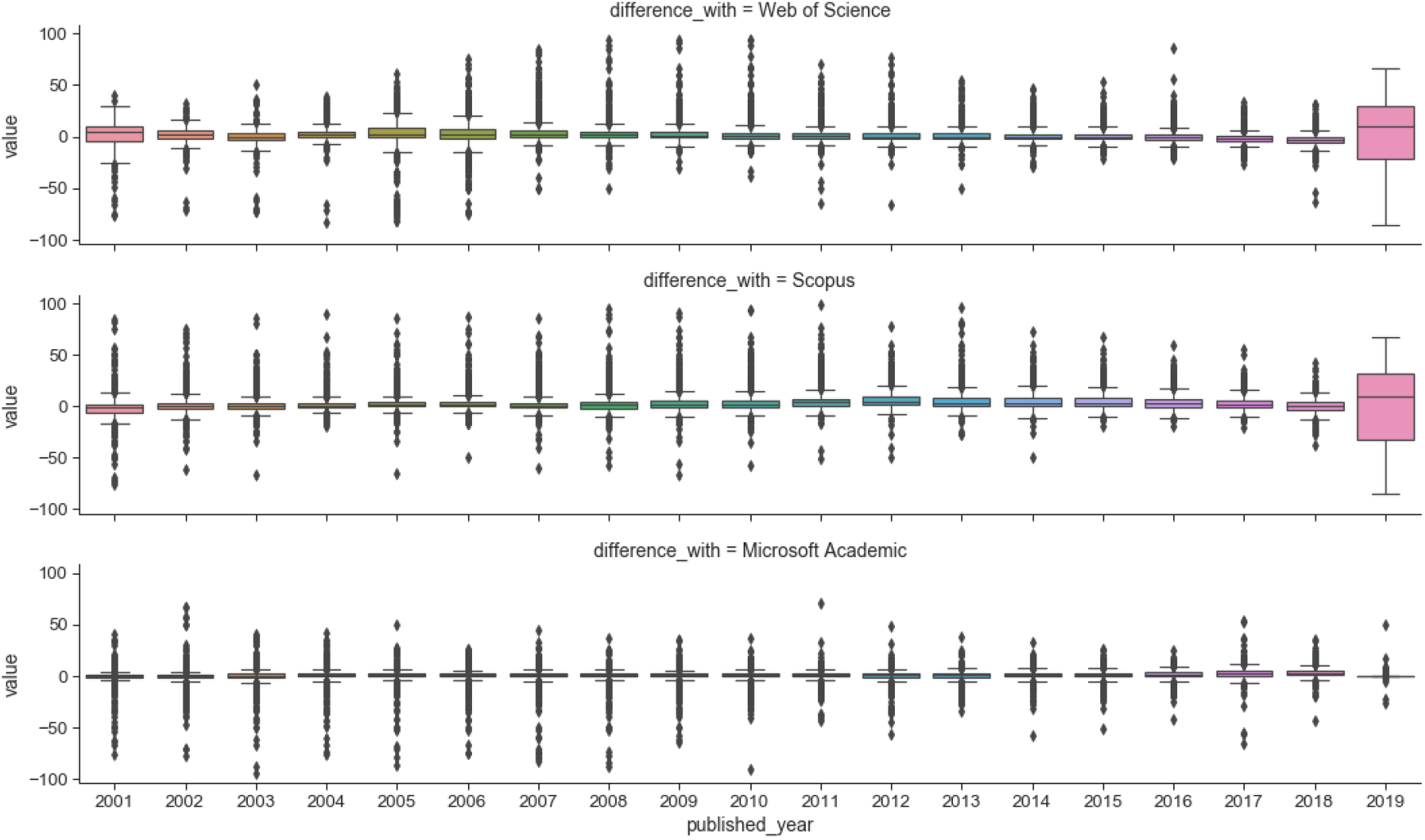
Difference in Total OA% over time between the full combined dataset against each of Web of Science, Scopus and Microsoft Academic.

**Figure 15:**
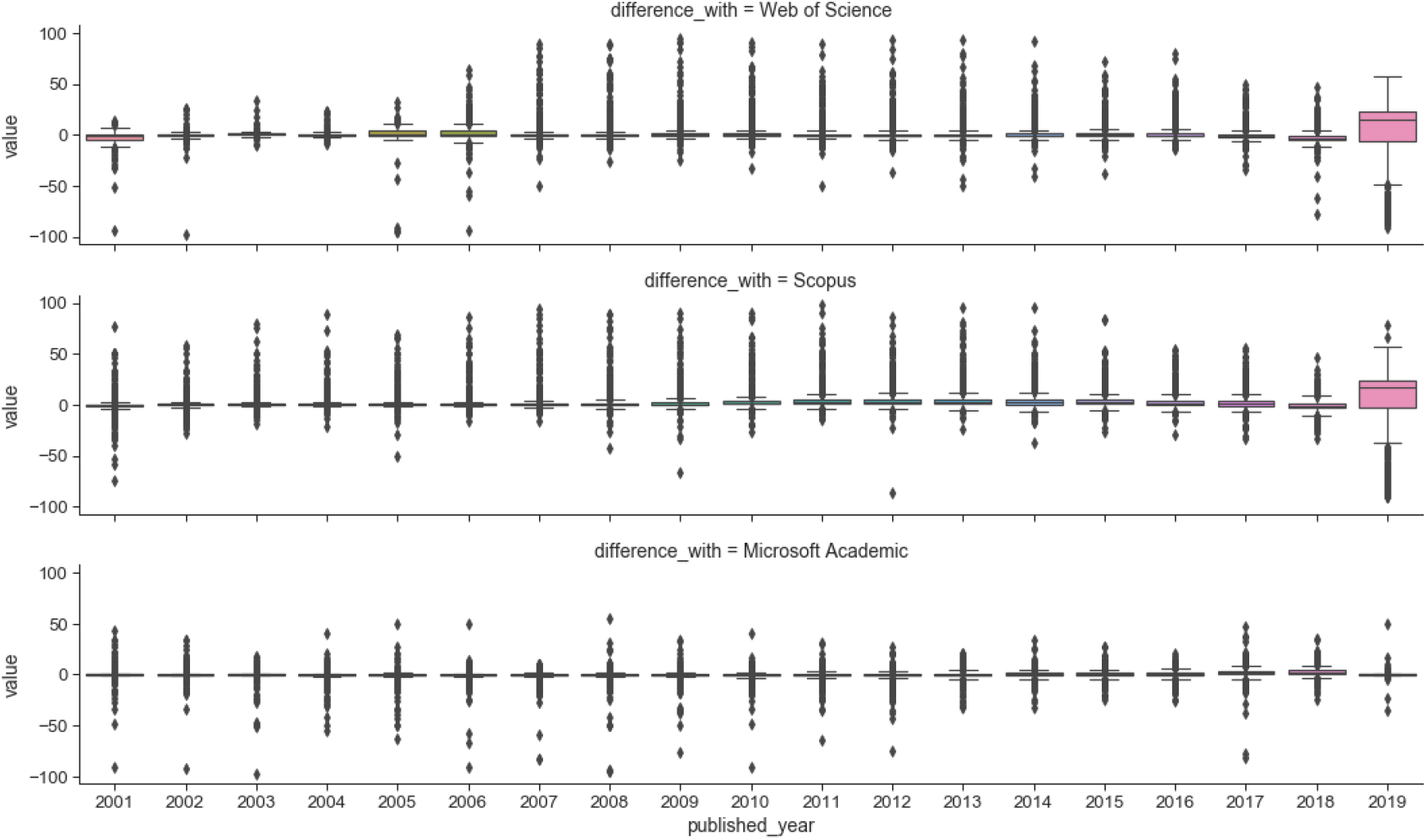
Difference in Gold OA% over time between the full combined dataset against each of Web of Science, Scopus and Microsoft Academic.

**Figure 16:**
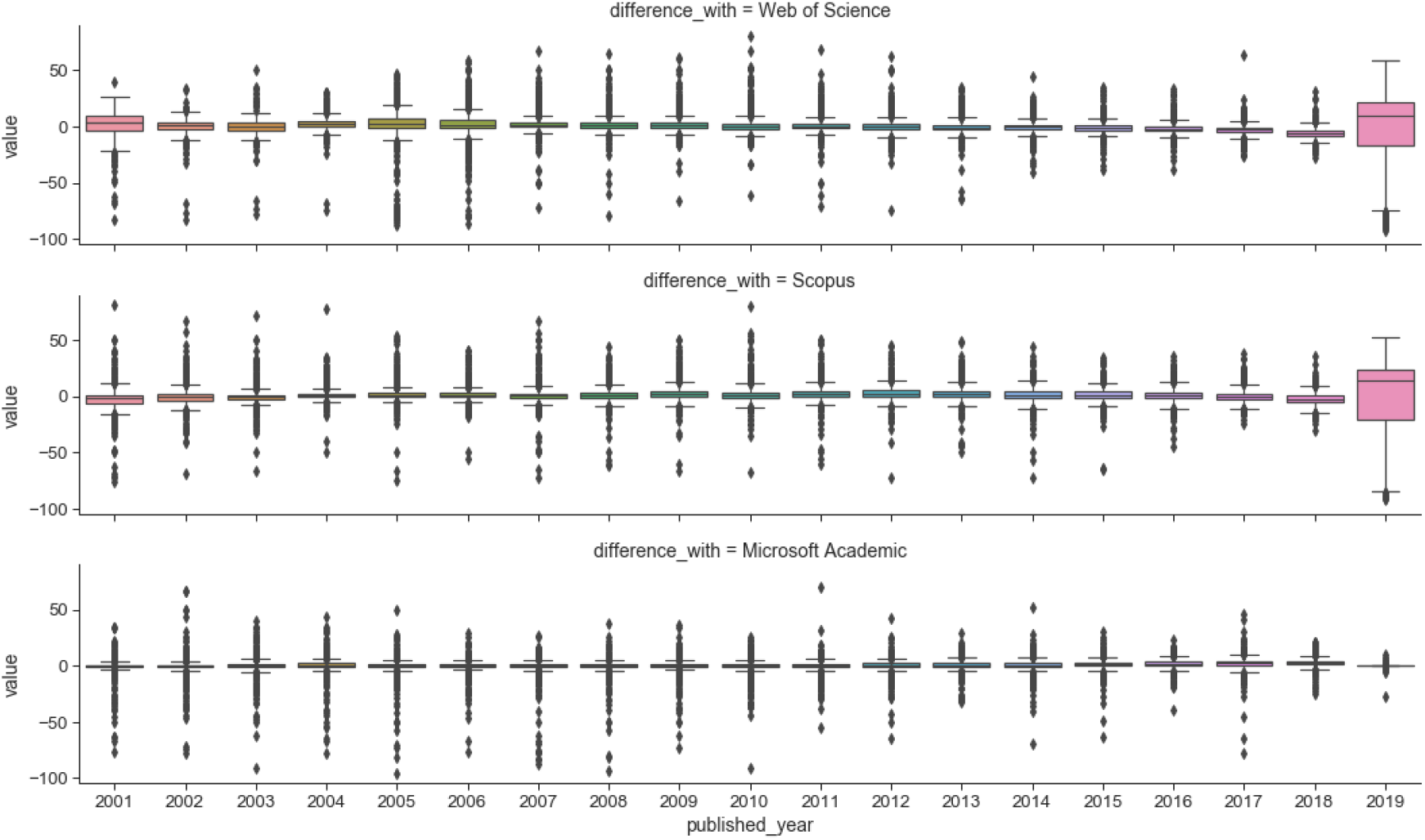
Difference in Green OA % over time between the full combined dataset against each of Web of Science, Scopus and Microsoft Academic.

The median difference across all boxplots is close to zero. However, a significant number of boxplots for the differences between the combined dataset against Web of Science and Scopus are characterised by positive skewness. This is particularly evidenced for the gold open access percentages, and followed by the total open access percentages. This implies a significant portion of the universities are assigned higher gold open access percentages by the combined dataset, as compared to using only Web of Science or Scopus. This seems to then result in higher total open access percentages as well. In contrast, the boxplots related to Microsoft Academic seem more symmetric and also seem to have a smaller spread (in terms of the interquartile range).

No clear trend is observed for comparisons across time. Except, there appears to be a slight increase in discrepancies (in terms of the total open access percentages and green open access percentages) as we move further back in time (evidenced by the increases in range and interquartile range).

Figure 17 presents the corresponidng comparisons of total open access percentages over time for pairs of Web of Science, Scopus and Microsoft Academic. Many of the boxplots for the difference between Microsoft Academic against the other two sources depicted positive skewness. There is again signals of very slight increase of differences further back in time.

**Figure 17:**
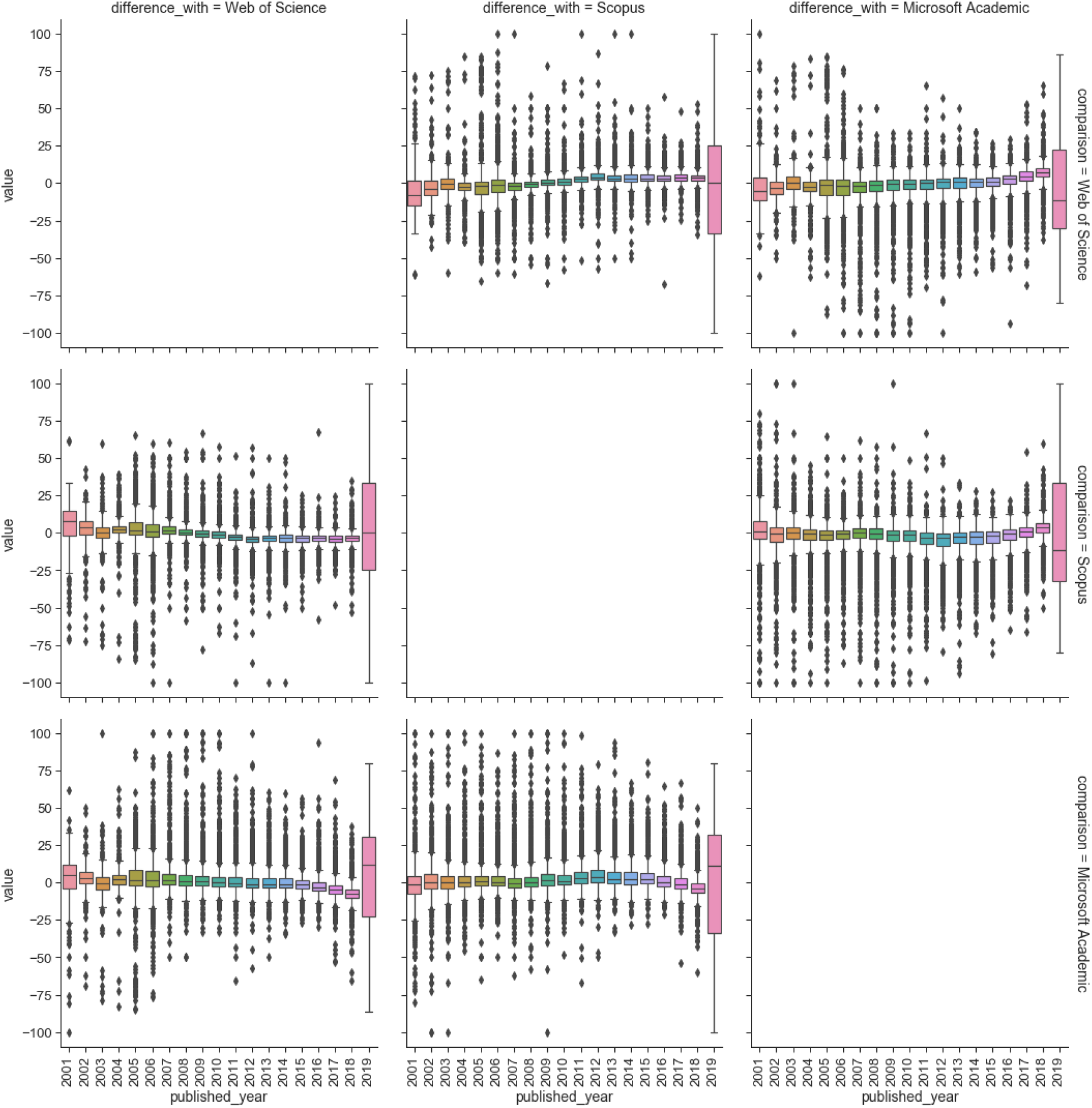
Difference in total OA% over time between any pairs of Web of Science, Scopus and Microsoft Academic.

Figure 18 compares the counts in total publication and various open access categories obtained throuh the combined datset against those derived from only individual sources. We first observe that alomst all dots lie below the diagonal line (few sit on the line), indicating that the combined dataset gives a significantly high number of counts in all categories as we would expect. For gold open access counts, we observe that Latin Amrican universities lie much further away from the diagonal line in the comparison against Scopus and Web of Science. This is an indication of the effect of the inclusion of Microsoft Academic have on universities from this region. It is also interesting to note that the number of outputs in the green open access and green in home reposities are dominated by European and North American universities.

**Figure 18:**
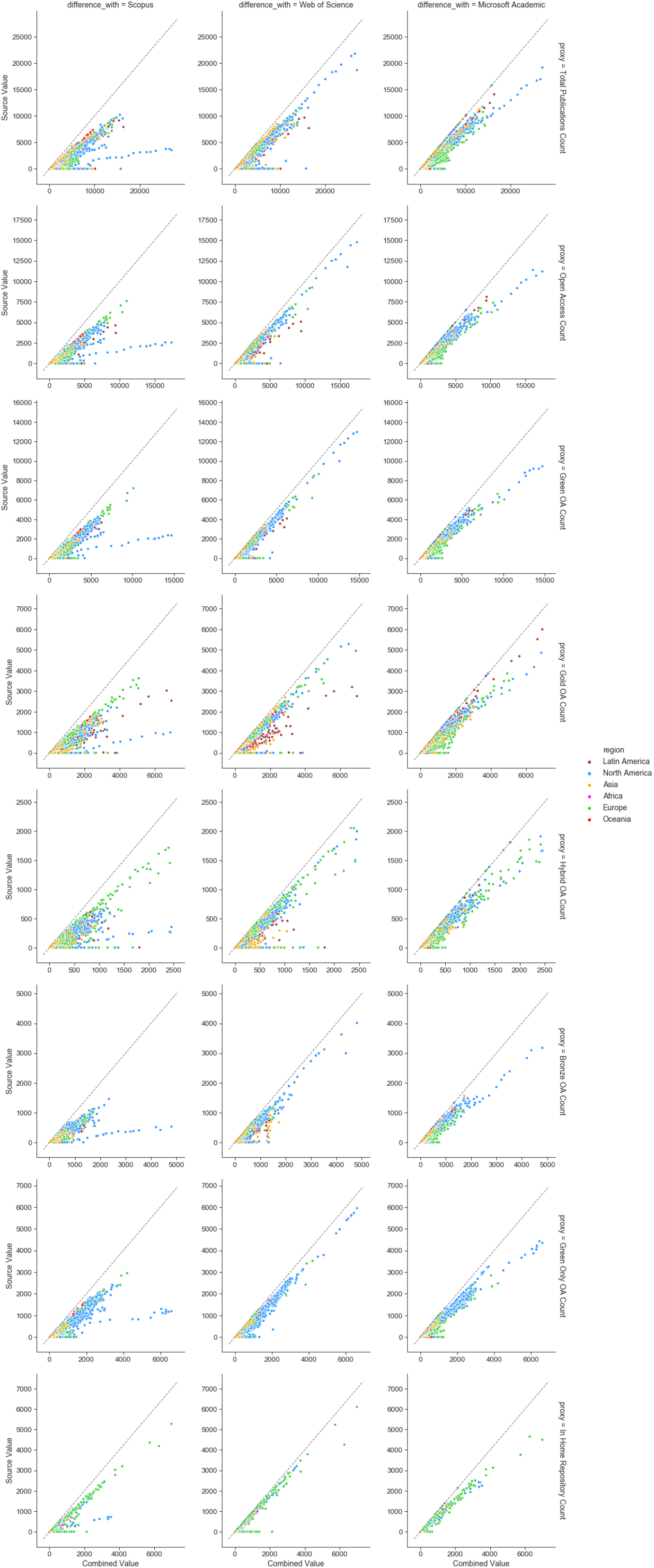
Scatterplots of the combined dataset against Web of Science, Scopus and Microsoft Academic, in terms of publication counts (in total publication, total OA, gold OA, green OA, hybrid OA, bronze OA, green only OA, and green in home repo OA) for all years in the dataframe.

In our data workflow, Scopus consistently give Harvard a lower number of outputs. This is shown by the slowly increasing line (of blue dots) on the bottom right of the plots in the first column above. This is due to the affiliation identifier in Scopus, where Harvard has multiple identifiers. The identifier in our database points to “Harvard University” but that does not cover all of “Harvard Medical School” which has a higher number of total outputs. This emphasises the importance of integrating multiple data sources to address differences between data sources.

Figure 19 presents corresponding scatterplots in terms of the difference categories of open access percentages. Several interesting patterns arise that highlights regional differences across the data sources. For the total open access percentages, we see almost a clear divide between European and North American universities, and the rest of the world (across the diagonal line), in the comparison of the combined dataset against Scopus and Web of Science. This is a clear evidence for the bias driven by Web of Science and Scopus. This is further explained in detail by observing the detailed open access categories. For example, the gold open access percentages for Latin American and Asian universities are significantly higher when using the combined dataset, in contrast to using only Web of Science or Scopus. Similarly, Asian universities are assigned significantly higher number of bronze open access through the combined dataset.

**Figure 19:**
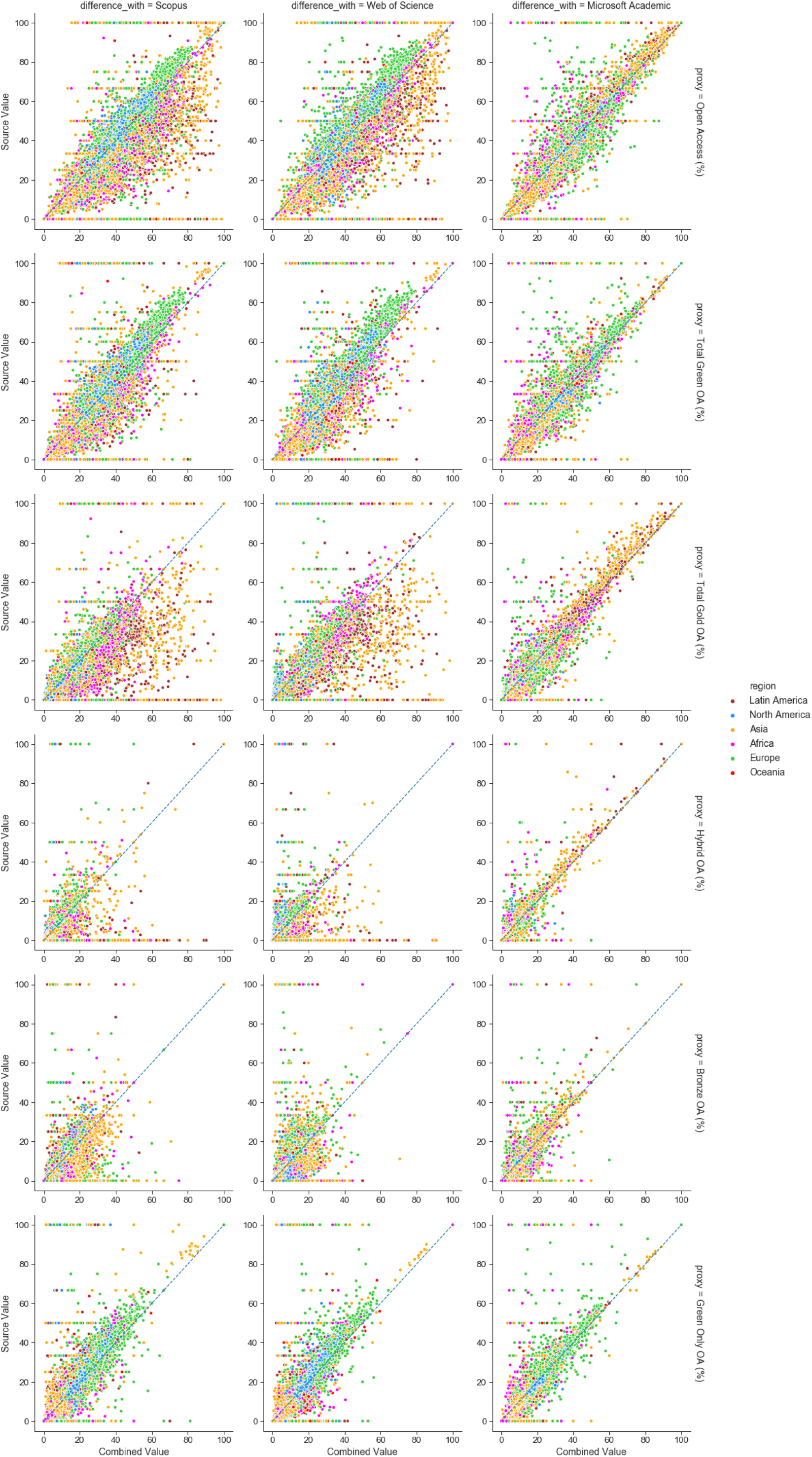
Scatterplots of the combined dataset against Web of Science, Scopus and Microsoft Academic, in terms of OA percentages (in total OA, gold OA, green OA, hybrid OA, bronze OA, and green only OA) for all years in the dataframe.

## 3 Sensitivity analysis on the use of different Unpaywall snapshot versions

In this section, we proceed to analyse the levels of sensitivity asscociated with the use of different Unpaywall data dumps. For these analyses, we maintain the use of the combined dataset, but use Unpaywall data dumps from different dates to determine open access status.

Figures 20, 21 and 22, compares our latest Unpaywall data dump against earlier versions for calculating total open access percentages, gold open access percentage, and green open access percentages, respectively, for 2017. Figure 20 shows that there are clearly more differences as we move towards earlier versions of Unpaywall, with many universities moving further away from the diagonal. However, Figures 21 and 22 further highlight that these differences are significantly due to the changes in green open access levels. This is an indication of Unpaywalll’s improving ability to capture data recorded by research repositories.

**Figure 20:**
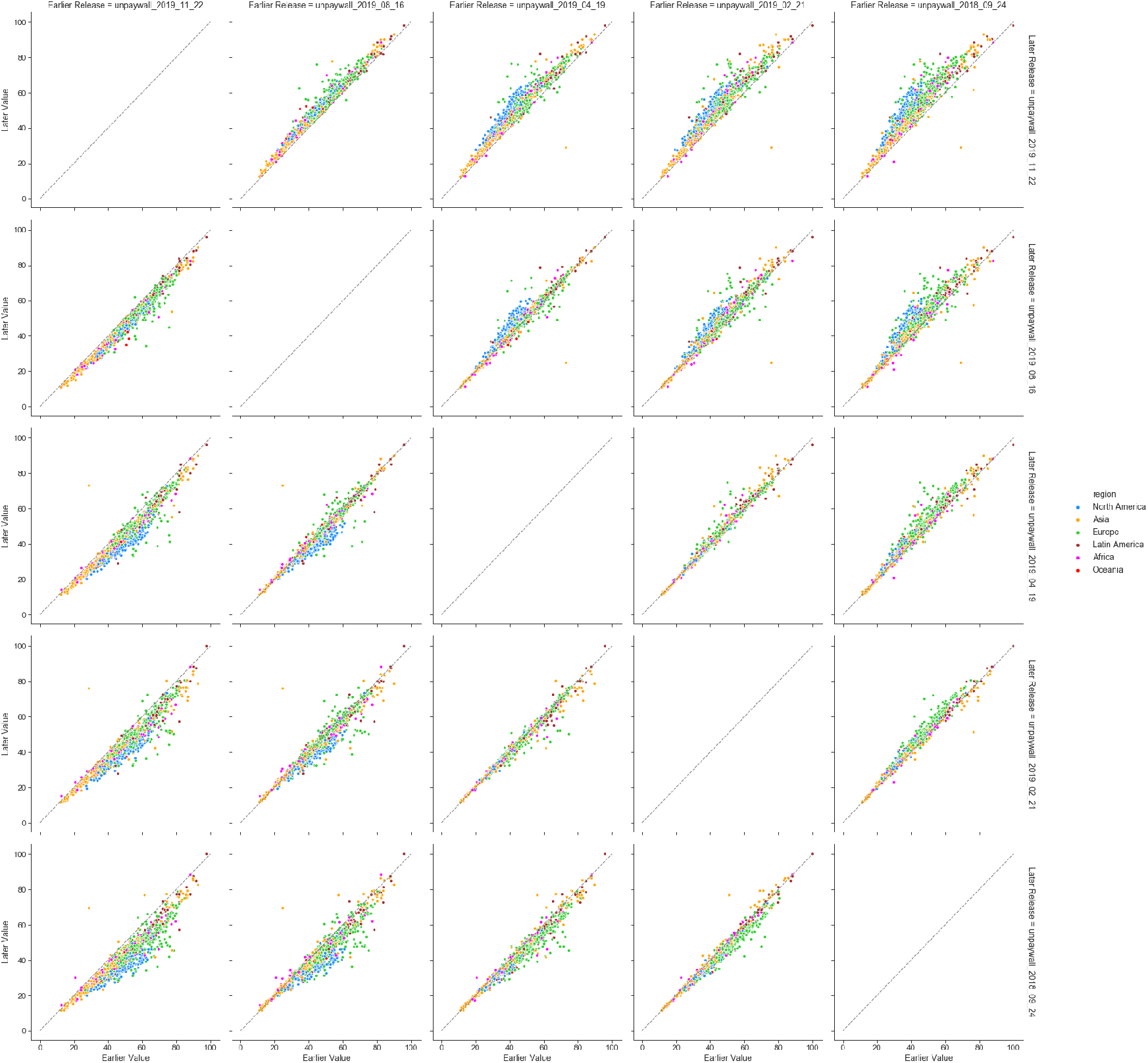
Scatterplots contrasting total OA% between various versions of Unpaywall data dumps.

**Figure 21:**
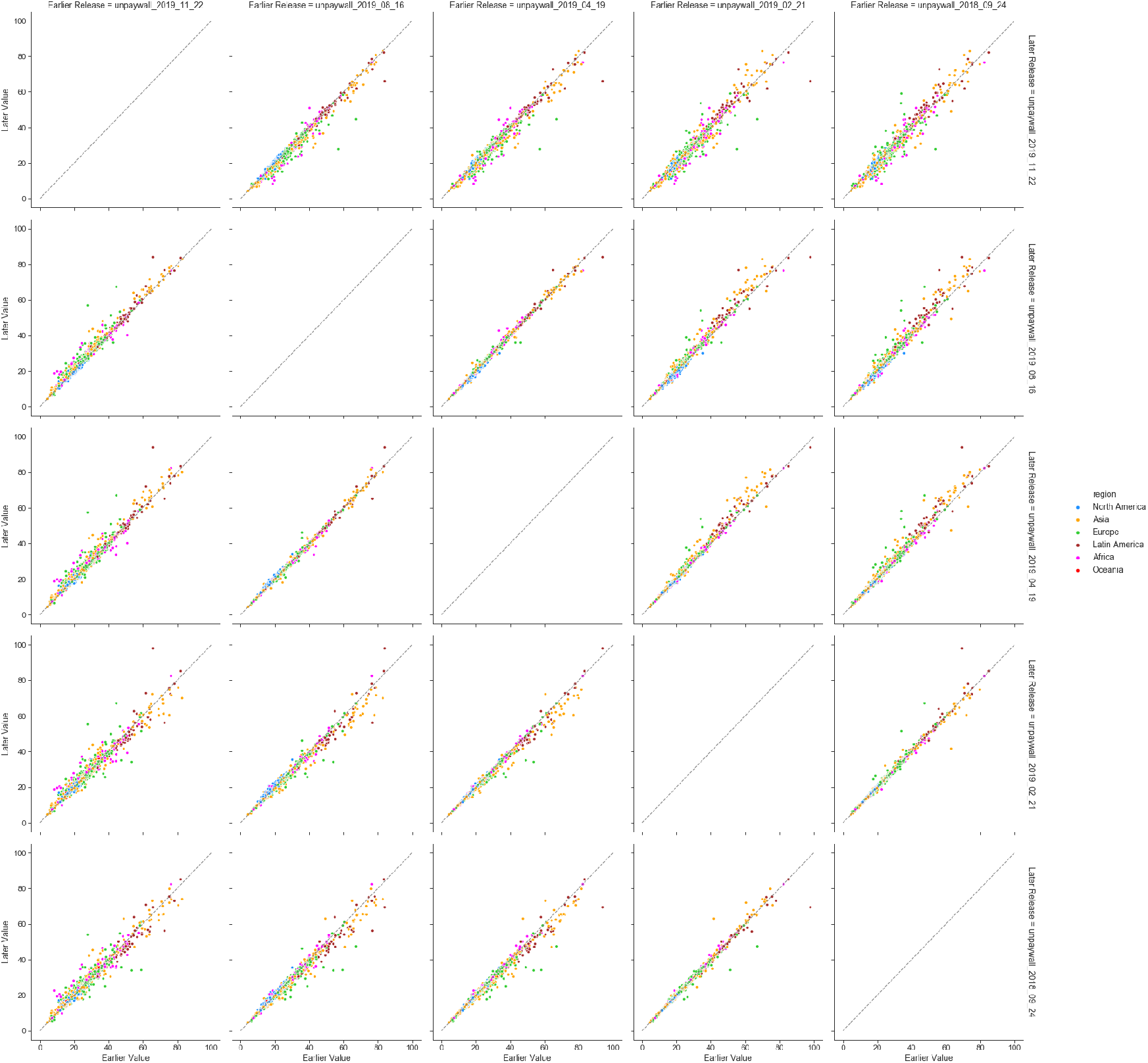
Scatterplots contrasting gold OA% between various versions of Unpaywall data dumps.

**Figure 22:**
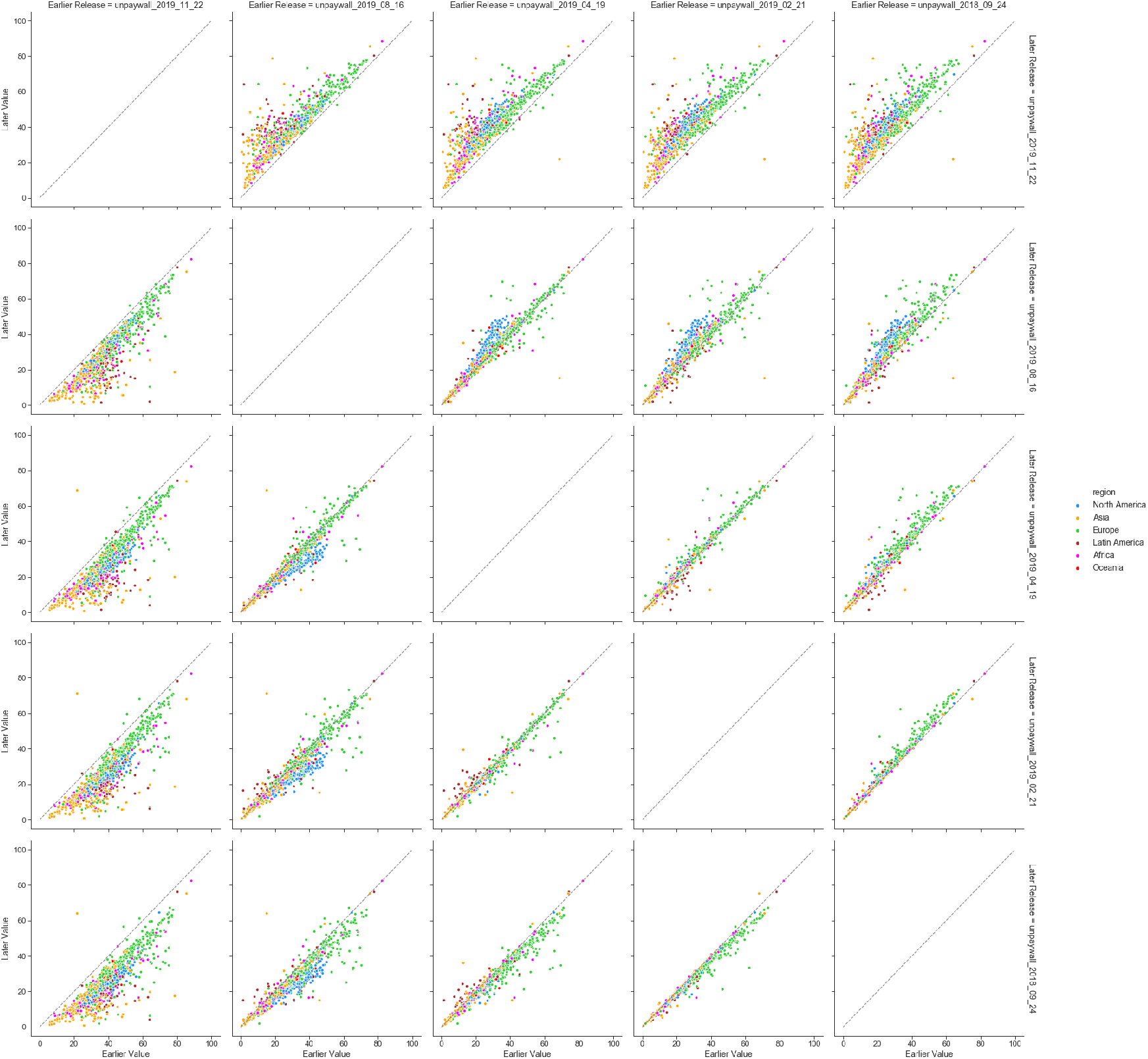
Scatterplots contrasting green OA% between various versions of Unpaywall data dumps.

Figures 23 to 27 depict the differences in total, gold, green, hybrid and bronze open access percentages (respectively) between the latest version of Unpaywall and several earlier versions. As observed earlier, larger differences are observed for total open access and green open access, as opposed to other open access categories.

**Figure 23:**
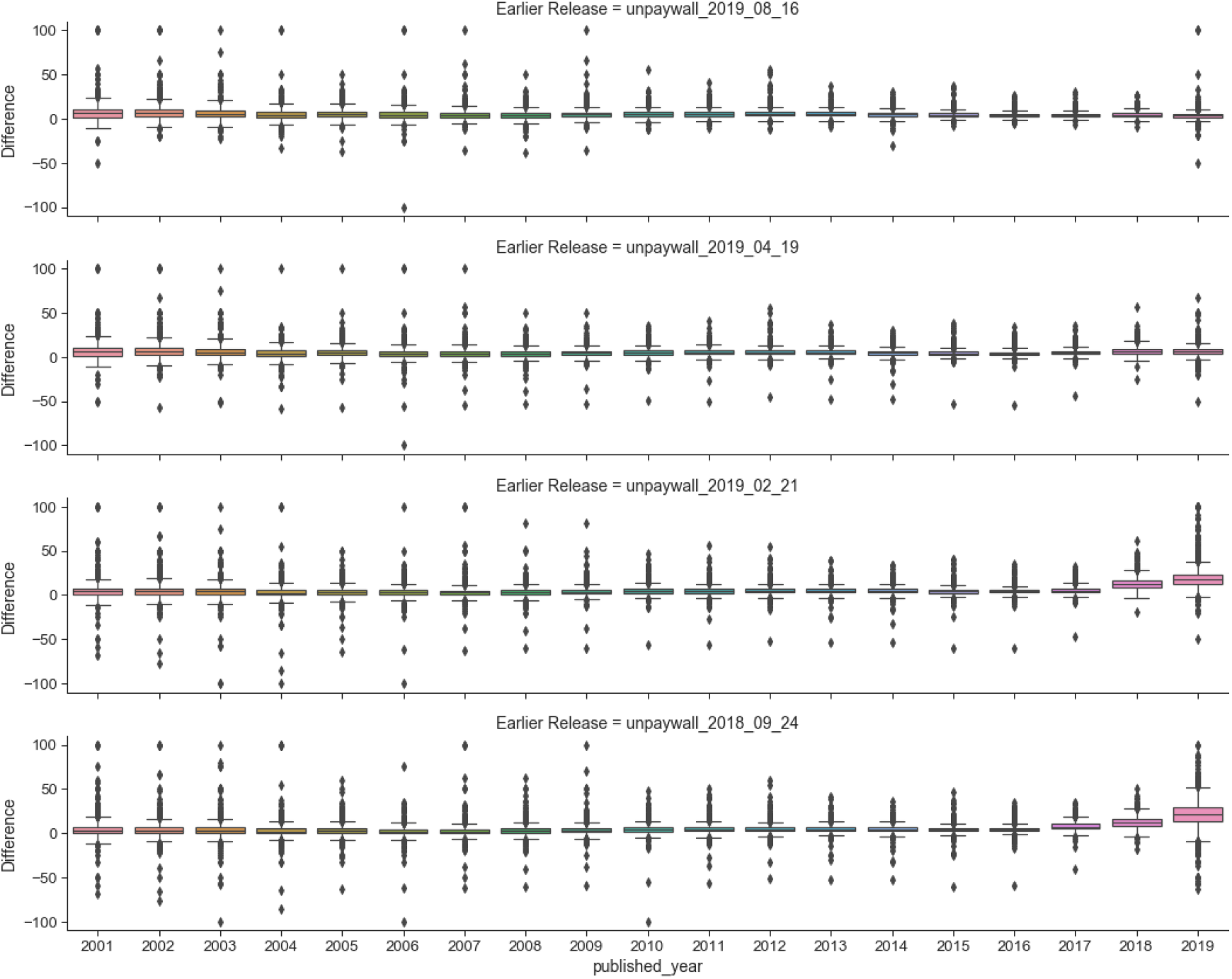
Comparing the most recent version of Unpaywall against earlier versions in terms of total OA%, for all years.

**Figure 24:**
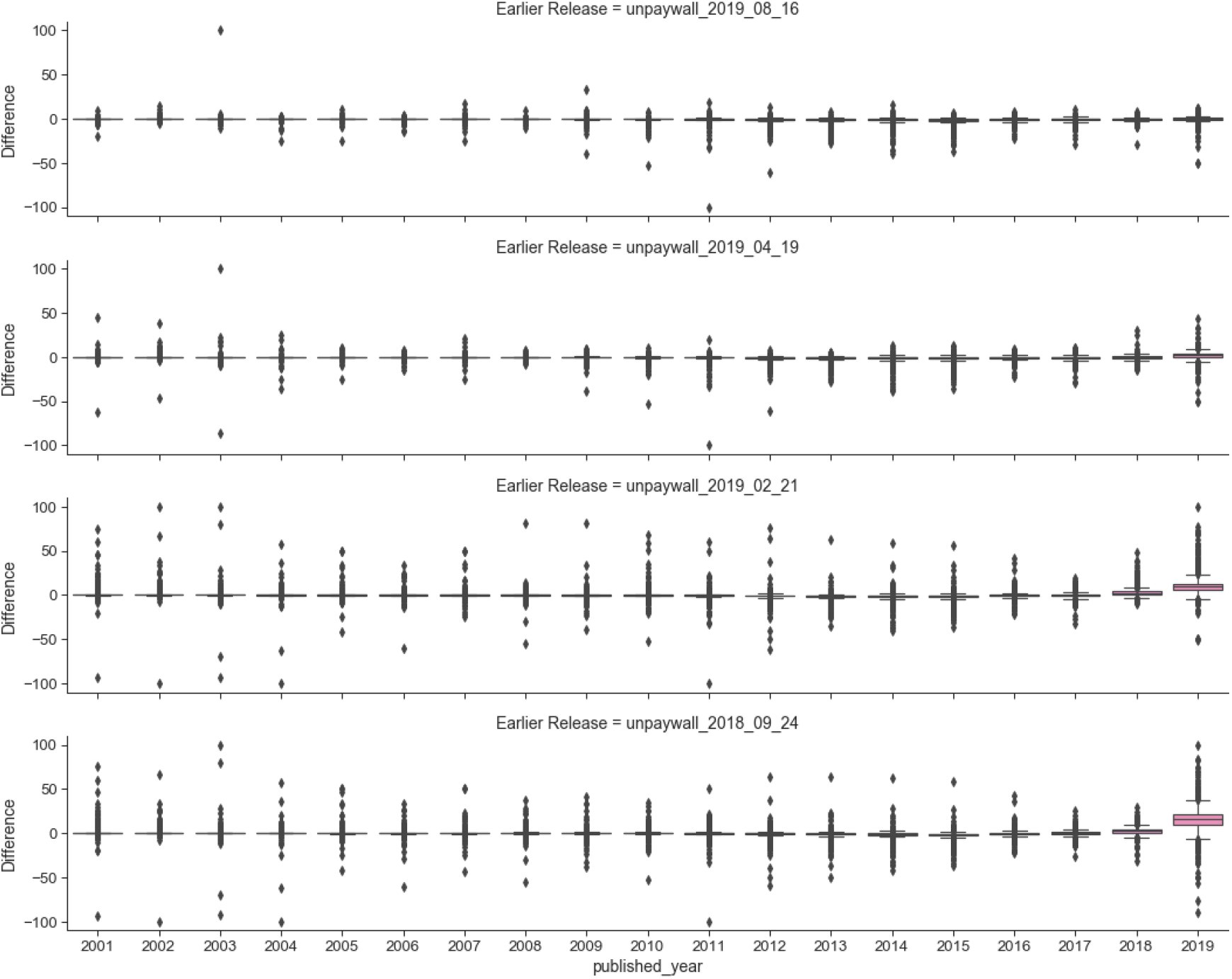
Comparing the most recent version of Unpaywall against earlier versions in terms of gold OA%, for all years.

**Figure 25:**
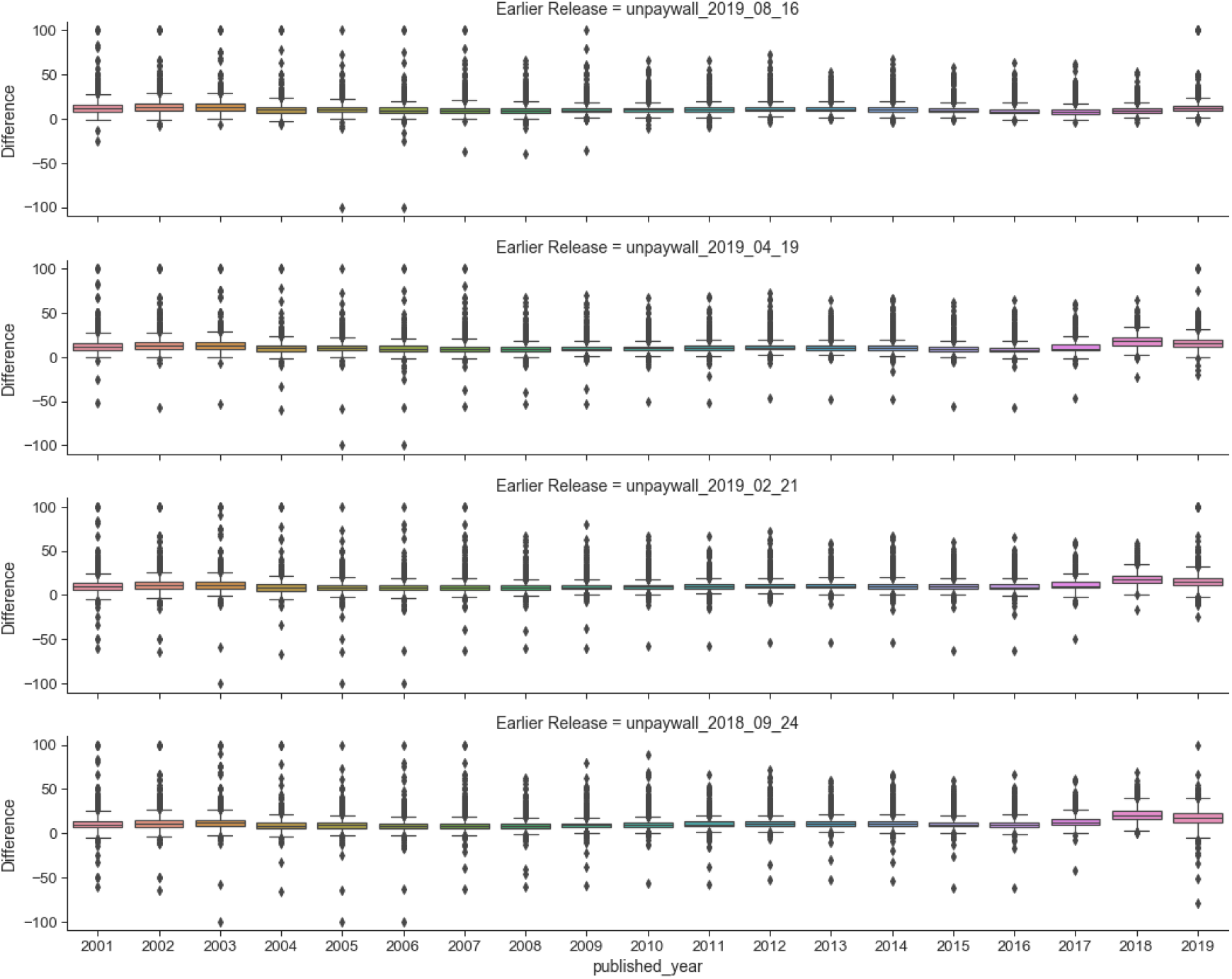
Comparing the most recent version of Unpaywall against earlier versions in terms of green OA%, for all years.

**Figure 26:**
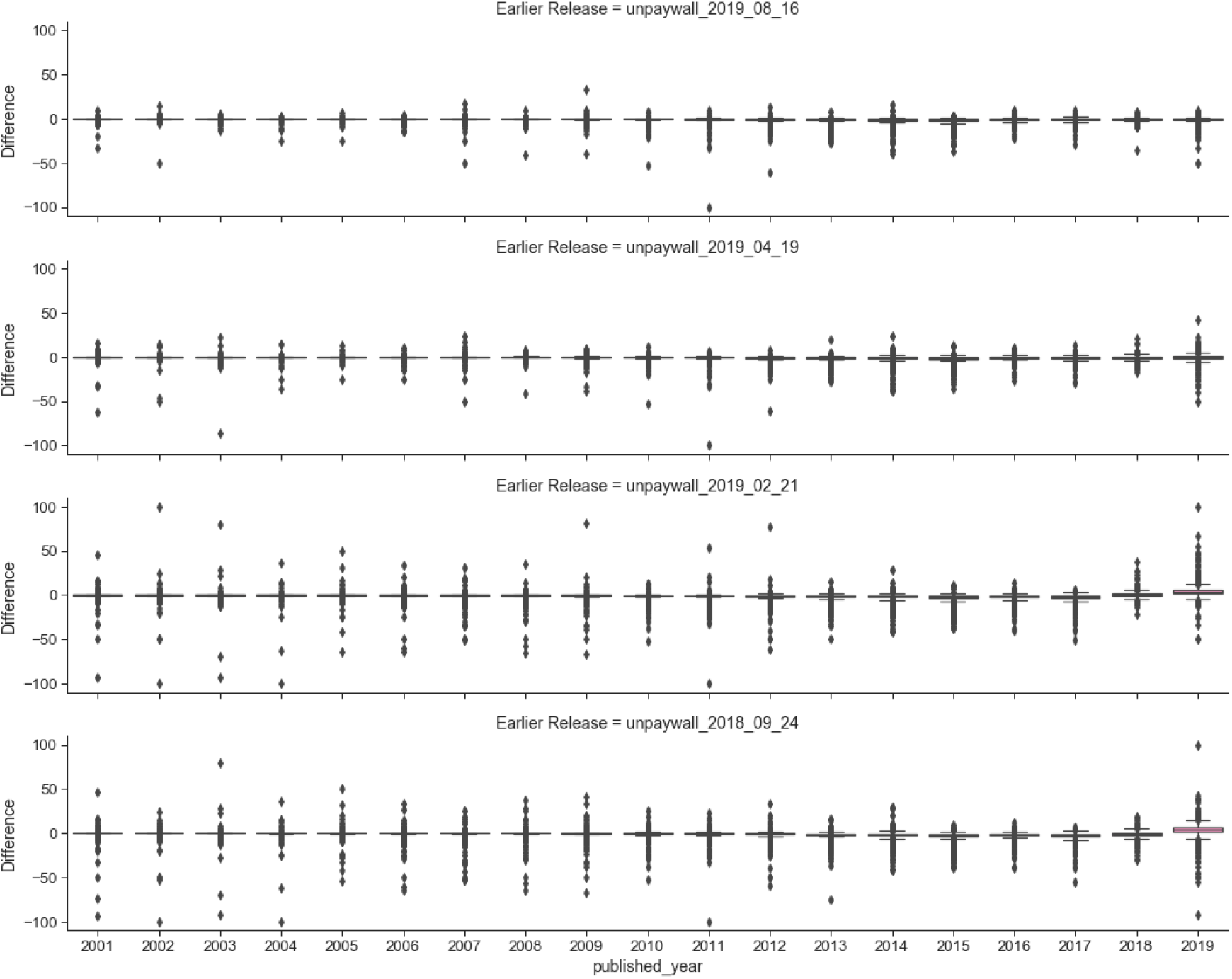
Comparing the most recent version of Unpaywall against earlier versions in terms of hybrid OA%, for all years.

**Figure 27:**
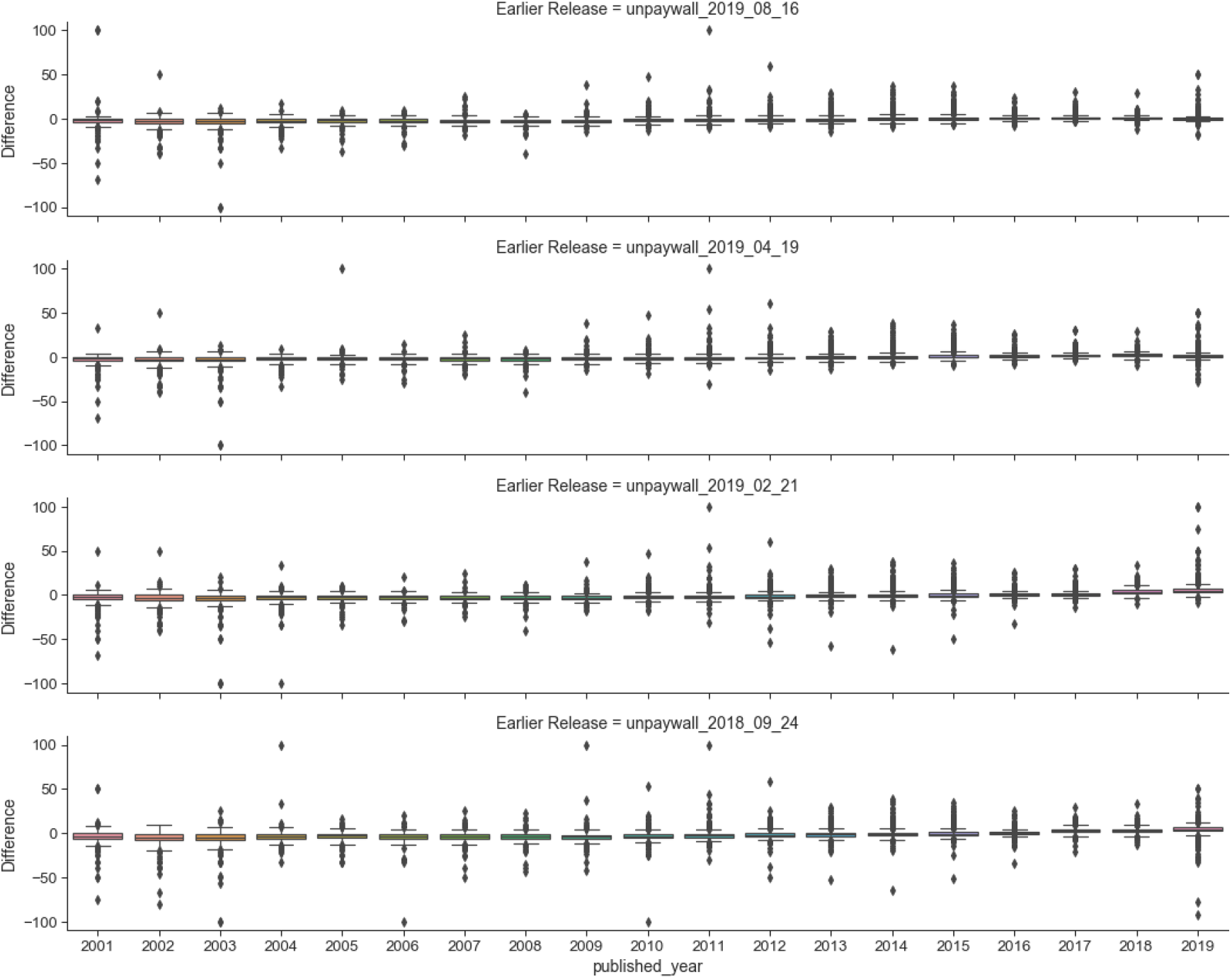
Comparing the most recent version of Unpaywall against earlier versions in terms of bronze OA%, for all years.

Two other interesting phenomenon are evidenced. Firstly, Unpaywall seems to be capturing a backfilling of historical data in particularly through the green and bronze routes. Secondly, we seem to observe the effects of embargos on self-archiving through the sudden jumps in 2018 for green open access, as we move towards earlier data dumps. To a lesser degree, some evidence of publisher embargos are shown in the gold open access.

## 4 Sensitivity analysis on sample size, margin of error and confidence levels

In this section we focus on analysing the sample size, margin of error and confidence levels associated with each university’s open access percentage estimates. In the corresponding main article (Huang et al, 2020b), margins of error were used to exclude universities that we have very little confidence for in terms of their open access levels produced through the data workflow. That means universities with a margin of level below a certain cut-off were not included in the list of Top 100 and some of the subsequent analysis. This is in addition to excluding universities that do not satify the conventional conditions for normal approximation to proportions (i.e., *np* > 5 and *n*(1 − *p*) > 5, where *n* is the sample size and *p* is the sample proportion for success).

For the choice of a cut-off level, we analysed the margins of error (equivalently, half of the length of the corresponding confidence intervals) at a number of confidence levels (i.e., 95%, 99% and 99.5%). In should be noted that we used the Šidák correction to control for the familywise error rate in multiple comparisons. This means that the multiple 95% confidence intervals calculated across the Top 100 universities are essentially 99.95% individual confidence intervals for each university. Hence, it justifies comparing the associated margins of error across a number of confidence levels in this range.

In Figures 28, 29 and 30, we plot the margins of error (at 95%, 99% and 99.95% respectively) against total, gold and green open access percentages, and against the total number of output, for each university. The aim is the spot a cut-off point for the margin of error when the data starts to behave more extremely. Subjectively, we consider the cut-off levels at 9.8, 12.8 and 17 for the three difference confidence levels. These correspond to approximately a sample size cut-off at 100 outputs, with a few exceptions. We consider this as a good starting point as we are focussed on research-intensive universities (Huang et al., 2020b) and we also would like a certain level of confidence about the open access estimates we provide. It is our aim to take a deeper exploration on issues around multiple comparisons (e.g., more advanced methods to cater for multiple comparisons) and principles for inclusion and exclusion.

**Figure 28:**
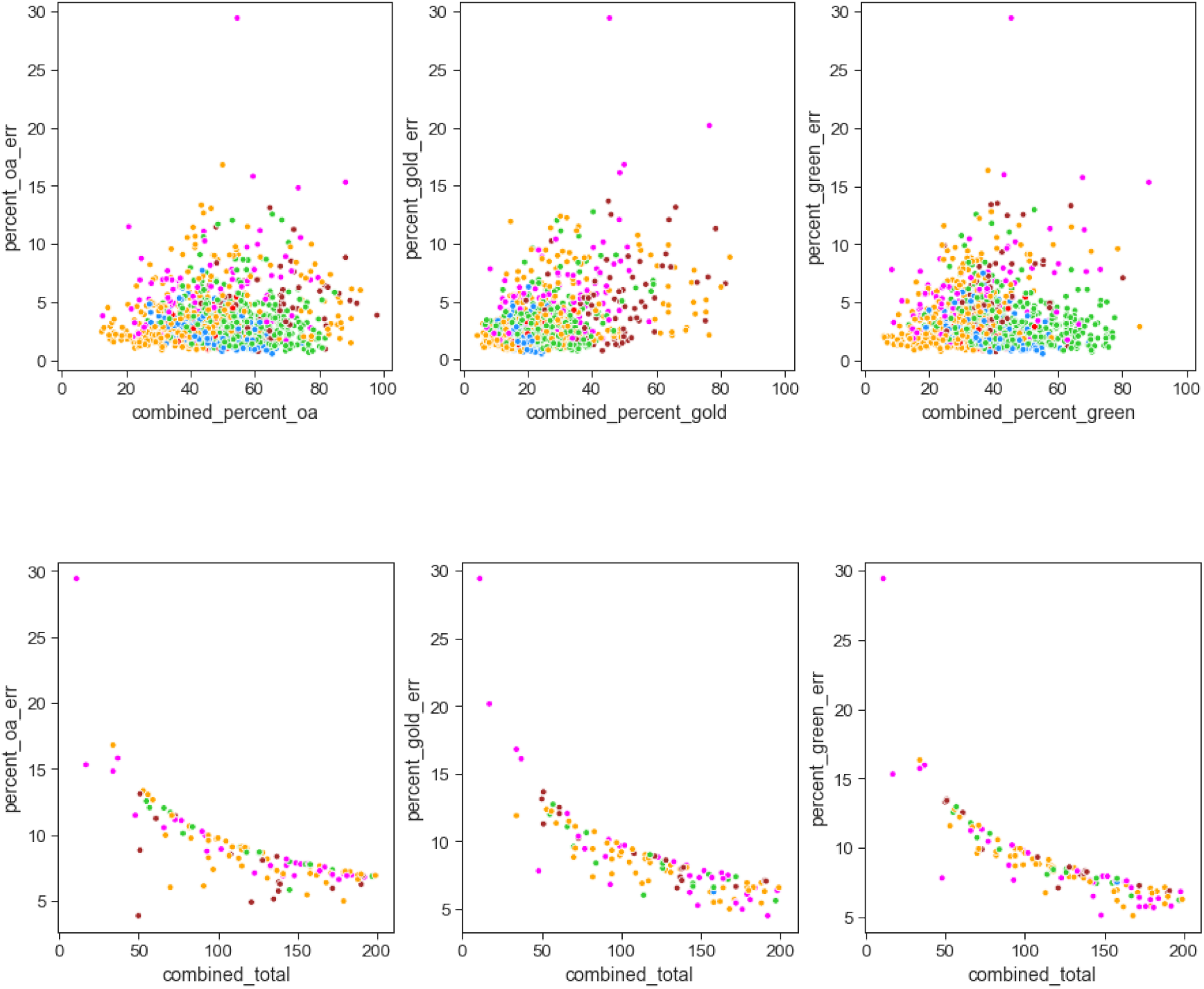
95% Margins of error versus open access percentages and the total number of publications for 2017.

**Figure 29:**
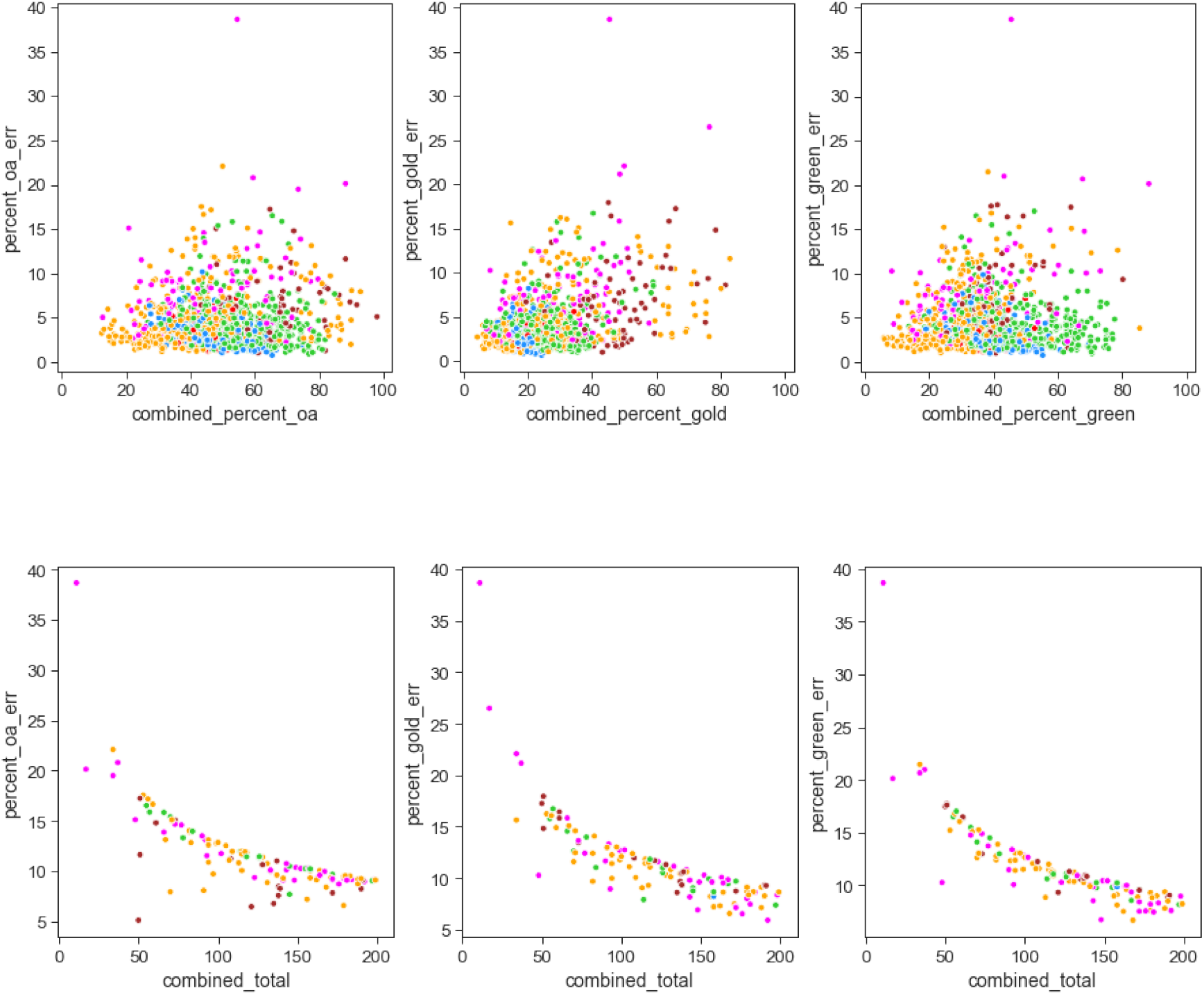
99% Margins of error versus open access percentages and the total number of publications for 2017.

**Figure 30:**
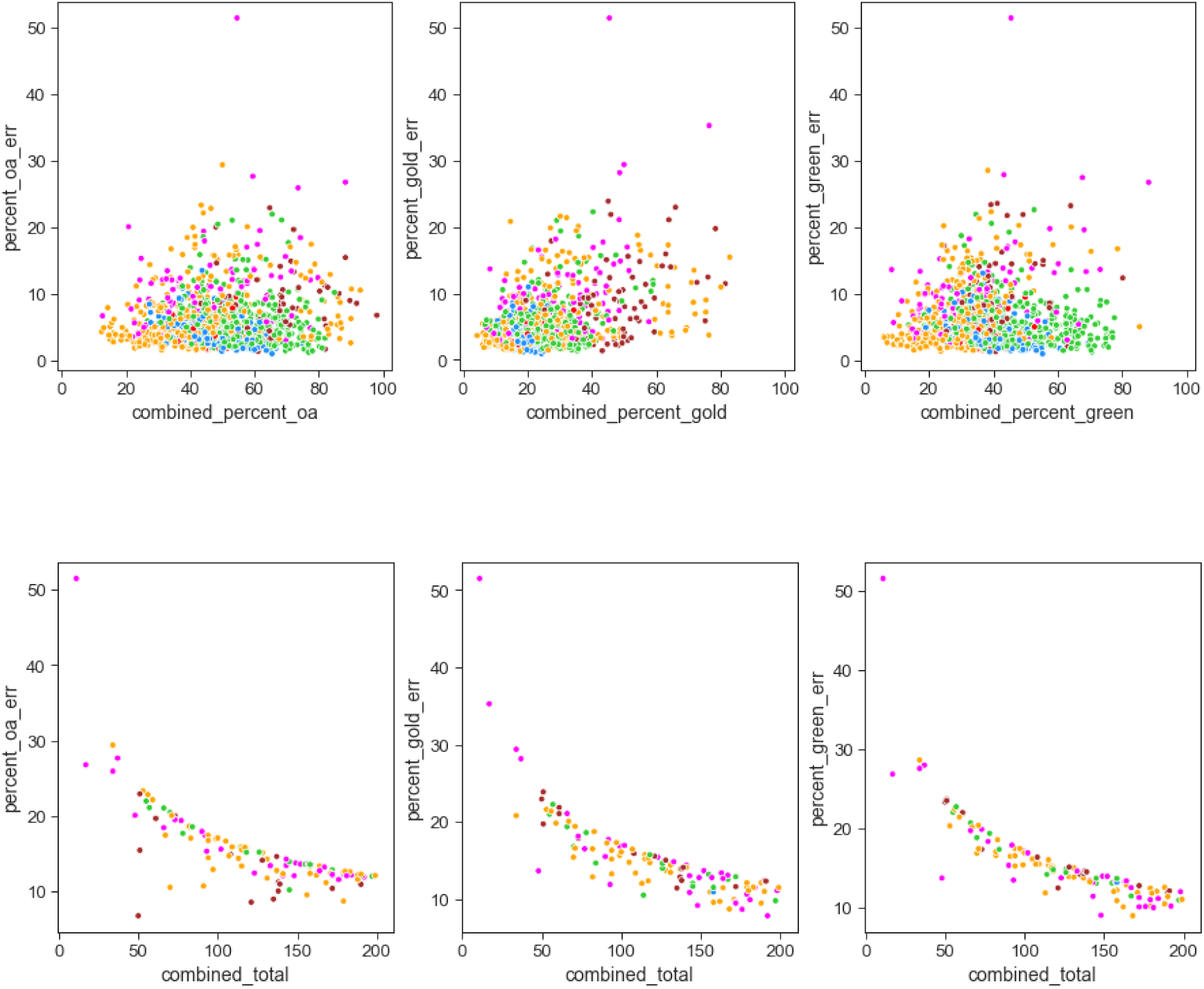
99.95% Margins of error versus open access percentages and the total number of publications for 2017.

## 5 Summary of main findings and implications

This article provided evidence for the sensitivities of comprehensive data workflow proposed in Huang et al. (2020b) against the choices of bibliographic data sources, versions of data sources, use of open access definitions, and confidence levels in statistical inference. Significant differences in the estimnated institutional open access levels can arise through choices made on these factors. These differences can also result in biases toward a university’s geographical location, preferred choice of open access route, and the time of publication.

This sensitivity analysis is essential for building a robust and fair evaluation framework for open access, and for understanding differences across data sources (in terms of coverage) and university groupings (in terms of regional foci). It also implies that any process for evaluating open access should clearly describe what data sources are used, which version of data is used, and how they are used in any standardisation and filtering procedures.

